# RNA binding protein IMP1/ZBP1 drives *Actb* mRNA localization and local translation in microglial peripheral processes and mediates morphological changes, motility and phagocytosis in response to inflammation

**DOI:** 10.1101/2023.07.31.551329

**Authors:** Josune Imaz-Iruretagoyena, Maite Blanco-Urrejola, Irene Núñez-García, Irene García-Toledo, Luis C Fernández-Beltrán, Mar Márquez, Silvia Corrochano, Amanda Sierra, Jimena Baleriola

## Abstract

Polarized cells in the brain, such as neurons and glia, rely on the asymmetric distribution of their proteins compartmentalizing the function of dendrites, axons, glial projections and endfeet. Subcellular proteomes can be assembled either by the transport of proteins synthesized in the cell soma or by the delivery of mRNAs to target compartments where they are locally translated into protein. This latter mechanism is known as local protein synthesis or local translation, and it has been best studied in neurons. Increasing evidence suggest it is also required to maintain local protein homeostasis in glial cells, however, in microglia, local translation remains largely unexplored. Given the scant evidence, we aimed at exploring the existence of local translation in peripheral microglial processes (PeMPs) and unravel its functional significance in response to inflammation, a major hallmark of neurodegenerative diseases. We report that local translation in PeMPs is enhanced by triggering a microglial inflammatory response with bacterial lipopolysaccharides (LPS). We found that *Actb* mRNA polarizes to PeMPs and is locally translated upon LPS exposure. mRNA localization in eukaryotic cells is driven by RNA binding proteins. Interestingly, downregulation of the *Actb* binding protein IMP1/ZBP1 impaired *Actb* mRNA polarization and its localized translation, and led to defects in filopodia distribution, PeMP motility, lamellar directed migration and phagocytosis in microglia. Thus, our work contributes to recent findings that mRNA localization and localized translation occur in microglia and gives a mechanistic insight into the relevance of this molecular mechanism in fundamental microglial functions in response to inflammation.

## INTRODUCTION

Brain cells present an extremely complex and polarized morphology with cell extensions (dendrites, axons and glial peripheral processes) reaching more than a meter in length in the case of some human axons ^1^ and more than 200 µm in some rat microglial populations ^2^. Such complexity is required for their specific functions, and it mirrors the equally complex structure of the organ they are hosted in. Morphological and functional polarization of brain cells is achieved by asymmetrical distribution of their proteins to distinct subcellular compartments. Subcellular proteomes are assembled either by protein transport from the soma or by local translation. The latter can be accomplished by the transport of the mRNAs to the periphery followed by their translation at the target site. This mechanism enables cells to react to environmental changes in an acute manner, as proteins are produced only when and where they are needed ^3^.

Although local translation is a highly conserved mechanism in eukaryotic cells ^4^, in the nervous system it has been mostly studied in neurons where it is involved in axon guidance, arborization, and maintenance ^5^, synapse formation ^6^ and synaptic plasticity^7^. Interestingly, in recent years, local protein synthesis dysregulation in neurons has been associated to many neurological disorders ^8^, and it has been attributed to RNA binding proteins (RBPs) malfunction ^9^. Local translation is not exclusive to neurons and increasing evidence indicate that oligodendrocytes, astrocytes and radial glia also rely on this mechanism to maintain local protein homeostasis and their functional fitness ^10^. Surprisingly, mRNA transport and localized translation in microglia have remained unexplored until only recently, when *Rpl4* mRNA, which encodes the 60S ribosomal protein L4, was detected in microglial processes in the mouse brain ^11^ and a localized ribosome-associated transcriptome was identified ^12^. Additionally, in an *ex vivo* wound model, microglial local translation was found to be relevant for efficient phagocytosis ^12^. However, localized translation in this cell type in the context of inflammatory insults has remained elusive. Neuroinflammation accompanies and/or is a recognized hallmark of neurodegenerative diseases ^13^. With this background, we wanted to determine if mRNA localization and localized translation in microglia were regulated by inflammation, and whether limiting the amount of *Actb* mRNA RBP IMP1/ZBP1 impaired microglial behaviour.

Microglia are the resident immune cells of the central nervous system, involved in brain homeostasis maintenance, and the first line of defence upon pathogen infection and brain injury. Microglia continuously survey the brain parenchyma with their peripheral processes which are continuously extending and retracting ^14^. Upon pathological insults, they rapidly change their morphology by remodelling their actin cytoskeleton, leading to enlarged processes and even the acquisition of bushy and ameboid morphologies ^15,16^. During neuroinflammation many secreted factors induce microglial proliferation and process migration towards the source of injury. Microglial motility depends on two peripheral F-actin-rich filamentous structures: 1) fine processes, termed filopodia, which survey the environment and respond to the inflammatory cues or chemokines by inducing intracellular signalling, and 2) large processes, termed lamellipodia, which are responsible for migration ^15,17^. Interestingly, these structures were already described in axonal growth cones more than three decades ago ^18^. Like in peripheral microglial processes (PeMPs), growth cone filopodia sense external cues and extend towards chemoattractants or collapse in response to molecular repellents guiding lamellae towards or away from the guidance cue. Importantly, growth cone behaviour is regulated by local protein synthesis ^5^. Given the similarities between axonal growth cones and PeMPs we hypothesized that mRNA transport and local translation happens in microglia supporting essential functional changes in response to inflammation.

Here we show that exposure of microglia to an inflammatory challenge driven by bacterial lipopolisaccharides (LPS) induces the localization of *Actb* mRNA to PeMPs both *in vitro* and *in vivo*, as well as its localized translation in primary microglial cultures. β-actin local synthesis has been involved in cell motility in fibroblasts ^19^, as well as in dendritic spine rearrangements and growth cone behaviour in neurons ^20,21^. One of the RNA binding proteins (RBPs) responsible for *Actb* localization is the well characterized IGF2 mRNA binding protein 1 / zipcode binding protein 1 (IMP1/ZBP1)^22^. Importantly, *Imp1* knockout mice show defects in *Actb*-containing granule motility in dendrites and transient *Imp1* knockdown affects the dendritic distribution of *Actb* and impairs dendritic arborization. Additionally, *Imp1* haploinsufficiency decreases *Actb* mRNA localization to axons and limits the regeneration of peripheral nerves upon injury ^23–25^. Interestingly, when we downregulated IMP1/ZBP1 in microglia, *Actb* localization and β-actin synthesis were decreased in PeMPs, and resulted in impaired filopodia distribution, PeMP motility, lamellar polarized migration and phagocytosis upon LPS exposure. Therefore, our results suggest for the first time that IMP1/ZBP1-dependent mRNA localization is relevant for the microglial response to inflammation.

## RESULTS

### LPS induces local protein synthesis in peripheral microglial processes (PeMPs)

To characterize RNA localization and local translation in microglia we first identified PeMPs, which we described as structures peripheral to the soma with stark accumulation of filamentous (F) actin and low or absent labelling of the endoplasmic reticulum-associated protein calreticulin (Figure 1A^i^). F-actin is virtually present in all cell types. Hence, we then wanted to determine if our primary cultures were enriched in microglia with few or no contamination from other cell types. We observed that 90.71 % (± 3.63 %) of all F-actin-positive cells (stained with phalloidin) expressed the microglia/macrophage marker Iba-1. Moreover, almost all Iba-1-expressing cells were also positive for the microglial marker P2Y12 (99.11 ± 0.25 %) (Figure S1A). Thus, by labelling cells with phalloidin we can more than likely specifically identify PeMPs. We then addressed if newly synthesized proteins could be detected in PeMPs by puromycin labelling. Puromycin is an aminoacyl-tRNA analogue that incorporates into nascent polypeptide chains during elongation in a ribosome-catalyzed reaction ^26^, and specific anti-puromycin antibodies can be used to detect *de novo* protein synthesis. We exposed cultured microglia to 2 µM puromycin for 10 and 30 minutes. Nascent peptides were readily detected after a 10-minute single exposure to puromycin both in the cell soma and in lamellar PeMPs, and signal increased with a 30-minute puromycin pulse. Cells preincubated with 40 µM of the protein synthesis inhibitor anisomycin showed reduced puromycin labelling (Figure S1B) confirming the suitability of puromycilation assays to detect newly synthesized proteins in the periphery of microglia.

**Figure 1.**
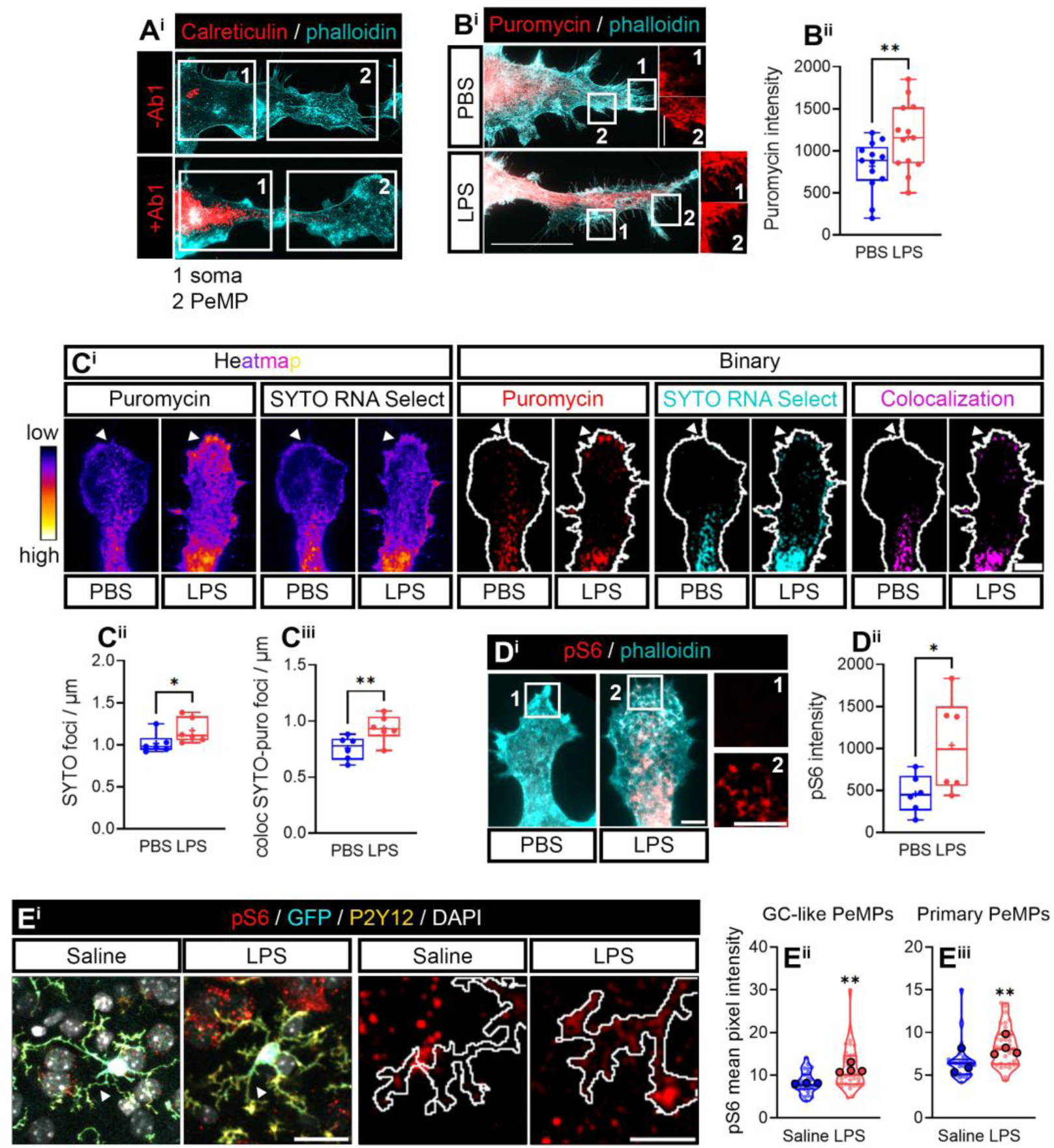
Newly-synthesized proteins increase in microglial peripheral structures upon LPS exposure. **(A)** To identify the somatic domain and the peripheral structures in microglia, cells were stained with an anti-carleticulin (Calr) antibody (+Ab1) to visualize the “somatic” ER (soma, 1) and with phalloidin to visualize the actin dense lamellipodia (2). The periphery was defined as the region where Calr was undetectable compared to a no-primary antibody negative control (-Ab1) which coincides with an intense phalloidin staining. Representative micrographs of the distribution of Calr and phalloidin are shown. Scale bar, 20 µm. **(B)** Newly synthesized proteins were measured in the periphery of microglia treated with vehicle (PBS) or LPS for 24 hours. Puromycilated newly synthesized proteins in the periphery of phalloidin-stained microglia are shown. Insets (1 and 2) show puromycin labelling at the edge of lamellae where filopodia emerge. Scale bar, 20 µm (insets, 5µm) (B^i^). The bar graph indicates the mean fluorescence intensity of puromycin in the periphery of microglia in 11 independent cultures (n=11) analyzed by two-tailed t test. **p < 0.01. (B^ii^). **(C)** Puromicyn and SYTO-positive heatmaps are shown, as well puromycin and SYTO-positive foci in binarized images. Scale bar, 5 µm (C^i^). Graphs represent the average SYTO-positive foci (C^ii^) and SYTO-puromycin colocalization (C^iii^) in the periphery of PBS- and LPS-treated microglia from 6 independent experiments (n=6) analyzed by two-tailed t tests. *p < 0.05; **p < 0.01. **(D)** Active ribosomal protein Rsp6 was analyzed. Insets show the levels of pS6 in PeMPs from PBS (1)- and LPS (2)-treated cells. Scale bars 5 µm (D^i^). The bar graph indicates the mean fluorescence intensity of pS6 in PeMPs in the periphery of microglia in 6 independent cultures (n=6) analyzed by two-tailed t test. *p < 0.05 (D^ii^). **(E)** pS6 staining in cortical microglia of fms-EGFP 1-month old mice injected with saline or LPS. Arrowheads in left panels indicate the GL-like shown in left panels. Scale bars 20 µm (left panels), 10 µm (right panels) (E^i^). Violin plots represent the mean intensity of pS6 in 27-36 sampled GC-like PeMPs (E^ii^) or primary processes (E^iii^) from 3-4 mice (n=27-36; smaller dots. N=3-4; bigger dots). Statistical analyses were performed by two-tailed t tests. **p < 0.01: ***p < 0.001.

Microglia sense signals from their environment, including inflammatory cues, with their processes ^12,14^. This likely requires changes in the local proteome for cells to rapidly adapt to changes in their surroundings. Thus, we sought to address if exogenously inducing inflammation regulated the levels of newly synthesized proteins in PeMPs. To that end we exposed primary cultures to LPS, an endotoxin commonly used to induce inflammatory responses in microglia ^27^, and treated cells with puromycin for the last 30 mins. Puromycin fluorescence increased upon a 24-hour LPS exposure in PeMPs (Figure 1B). A pre-requisite for local protein synthesis is the transport of mRNAs to peripheral processes and the presence of ribosomal components. To determine if LPS-induced increase of puromycin in PeMPs was a result of local translation, we addressed the presence of RNA in PeMPs in control and LPS-stimulated microglia using SYTO RNASelect, a fluorescent dye that selectively binds RNA ^28^ showing both a diffuse and punctate staining (Figure S1C). We analyzed SYTO-positive foci and SYTO-puromycin colocalization, as previously described ^29^, in microglia exposed to vehicle (PBS) or LPS for 24 hours. LPS treatment increased SYTO foci (binarized images in Figure 1Ci and Figure 1Cii) in PeMPs as well as SYTO-puromycin colocalization, especially at the edge of lamellipodia where fine processes (filopodia) emerge (binarized images in Figure 1Ci and Figure 1Ciii). Additionally, we determined if the protein synthesis machinery was activated in LPS-treated cells. For this, we immunostained microglia with an antibody that recognizes the phosphorylated form of ribosomal protein Rsp6 (pS6). Rsp6 phosphorylation has been previously used as a redout for local protein synthesis in neuronal axons ^30,31^. Our results indicated a significant increase of pS6 in PeMPs from LPS-treated microglia compared to control cells (Figure 1D). Finally, to determine if local protein synthesis was enhanced in response to inflammation in a more physiologically relevant model, we intraperitoneally injected 1-month-old MacGreen mice (fms-EGFP) with saline or LPS for four consecutive days. The MacGreen strain expresses enhanced green fluorescent protein (EGFP, here GFP) under the promoter of colony stimulating factor 1 receptor (*Csf1r*) and enables the visualization of the macrophage/microglia lineage in the brain ^32^. We observed that in LPS-injected mice all GFP-positive cells in the cortex expressed the microglia-specific protein P2Y12 (data not shown) indicating no infiltration of border macrophages. We then focused on this microglial population to analyze the levels of Rps6 phosphorylation in PeMps *in vivo*. We observed that pS6 was significantly increased in response to LPS in PeMPs that resembled axonal growth cones (GC-like PeMPs. Figures 1Ei and 1E^ii^) as well as in primary processes that directly emerge from the cell soma (primary-PeMPs. Figure 1E^1^ and 1E^iii^). Altogether, these data strongly suggest that LPS-induced inflammation enhances local protein synthesis in PeMPs both *in vitro* and *in vivo*.

### LPS drives *Actb* mRNA localization to PeMPs

Like in the axonal growth cones, PeMPs’ fine processes (filopodia) sense external cues and extend towards chemoattractants or collapse in response to molecular repellents guiding large processes (lamellae) toward or away from the guidance cue. Importantly, growth cone behaviour is regulated by local protein synthesis. To identify candidate transcripts whose local translation could be potentially regulated in response to inflammation, we performed RNA sequencing of control PeMPs isolated in a transwell system which consists of a culture insert with a polyethylene terephthalate membrane that creates two compartments. Cells were seeded on top of a 1 µm-pore membrane, that enables microglial PeMP extension toward the lower compartment restricting cell body migration (Figure 2Ai). Total RNA was isolated from the lower (PeMps) and upper compartments (whole lysate). We then focused on RNAs present in PeMPs at qualitatively similar or higher levels than in whole cell lysates. Given the similarities between PeMPs and axonal growth cones, we searched for transcripts involved in growth cone behaviour in response to guidance cues to determine to what extent were these transcripts localized to PeMPs as well. We focused on three of those RNAs that were present our PeMP transcriptome dataset: *Par3*, *Actb* and *Rhoa* ^20,33,34^ (Figure 2Aiii). We performed fluorescent *in situ* hybridization (FISH) and observed that none of them could be detected above background levels (established by the FISH signal from non-targeting probes) in PeMPs after a 24-hour treatment with LPS (Figures S2A and S2C^iii.^ *RhoA* data not shown). However, *Actb* mRNA was readily detected in peripheral processes (Figures S2C^i^ and S2C^ii^) and increased in response to a 30-minute exposure to LPS (Figures B^i^ and B^ii^). *Par3*, on the other hand, could also be detected in PeMPs following short treatments (Figure S2C^i^ and S2C^ii^) but no changes were induced by acute inflammation (Figures 2Ci and 2C^ii^). FISH experiments for RhoA detection in PeMPs yielded no conclusive results (data not shown). Thus, we further focused on *Actb* and *Par3*. Frequency analysis obtained from binarized images indicated that *Actb* granules accumulated away from the soma inmicroglia after 30-minute LPS treatments but not in control cells (Figures 2Biii-2B^iv^). *Par3* foci, however, were significantly higher in PeMPs compared to the soma both in PBS- and LPS-treated cells (Figures 2Ciii-2C^iv^), suggesting basal *Par3* localization to the periphery of microglia. These results suggest that acute LPS exposure leads to the polarization of *Actb* mRNA toward PeMPs *in vitro*.

**Figure 2.**
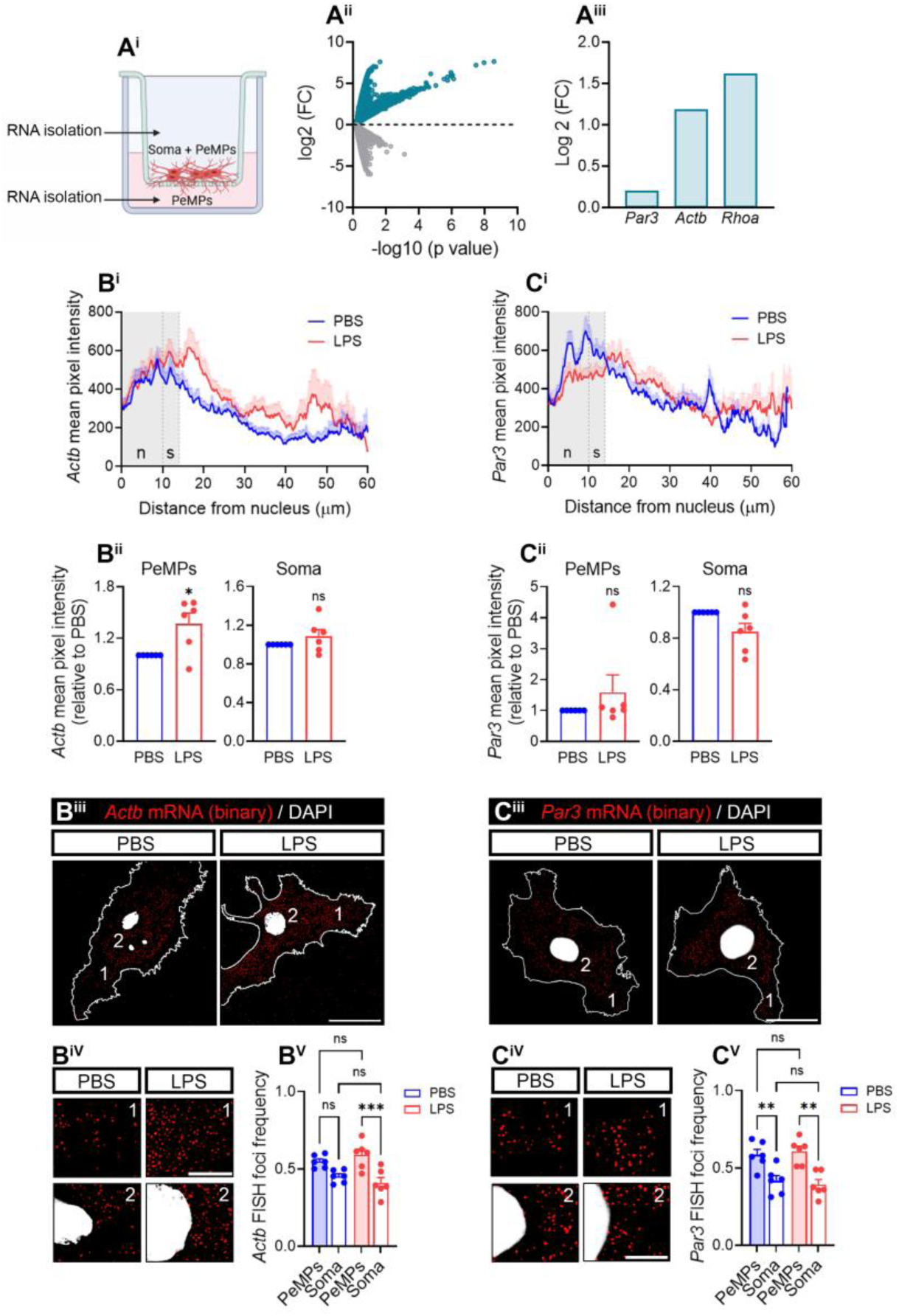
LPS increases *Actb* in PeMPs *in vitro*. **(A)** Schematic representation of a transwell with 1-µm-diameter pore used for the isolation of PeMPs and whole cell lysates. Microglia were seeded on of the membrane enabling microglial PeMP extension toward the lower compartment (A^i^). Total RNA was isolated from both compartments. Trancripts at similar or higher levels than in whole lysates are depicted in cyan (A^ii^) and three of these mRNAs (involved in axonal growth cone behaviour) are shown (A^iii^). **(B)** *Actb* levels measured by FISH in primary microglia treated with PBS in LPS for 30 minutes. Linescans represent the distribution of the FISH signal from the nucleus to the PeMPs in 45-48 individual cells per condition (B^i^). Bar graphs show *Actb* levels in LPS-treated cells relative to controls in PeMPs and the soma measured in 6 independent experiments (n=6). Two-tailed t tests. *p < 0.05; n.s, not significant (B^ii^). Distribution of *Actb*-positive foci in binarized images (B^iii^). (1) and (2) indicate the PePMs and the perinuclear regions respectively represented in insets Scale bars, 20 µm (10 µm insets) (B^iv^). The bar graph shows the relative frequency distribution of *Actb* foci in the periphery and the soma of microglia after treatment with PBS or LPS for 30 minutes in 6 independent experiments (n=6). One-way ANOVA followed by Holm-Šídák’s multiple comparison test. ***p < 0.001; ns, not significant (B^v^). **(C)** *Par3* levels measured by FISH in primary microglia treated with PBS in LPS for 30 minutes. Linescans represent the distribution of the FISH signal from the nucleus to the PeMPs in 49-58 individual cells per condition (C^i^). Bar graphs show *Par3* levels in LPS-treated cells relative to controls in PeMPs and the soma measured in 6 independent experiments (n=6). n.s, not significant (C^ii^). Distribution of *Par3*-positive foci in binarized images (B^iii^). (1) and (2) indicate the PePMs and the perinuclear regions respectively represented in insets Scale bars, 20 µm (10 µm insets) (C^iv^). The bar graph shows the relative frequency distribution of *Par3* foci in the periphery and the soma of microglia after treatment with PBS or LPS for 30 minutes in 6 independent experiments (n=6). One-way ANOVA followed by Holm-Šídák’s multiple comparison test. **p < 0.01; ns, not significant (B^v^).

We then asked whether these transcripts were regulated *in vivo*. We performed FISH in the cortex of 1-month-old fms-EGFP mice exposed to saline or LPS by IP injections and focused on the microglial population positive for both GFP and P2Y12 (Figure 2Ci and Figure S3Ci). As shown in the linescans, we observed a generalized increase of *Actb* in all cell compartments (Figure 3Ai) in response to LPS, including GC-like PeMPs and the soma (Figure 3Aii). We obtained similar results for *Par3* mRNA (Figures S3A-S3C). However, unlike *in vitro*, *Actb* and *Par3* polarization were unclear as the intensity from the FISH signal was higher in the soma than in PeMPs both in saline- and LPS-injected mice. Despite these discrepancies, which are likely a result on differences in the experimental approaches, we concluded that at least in the case of *Actb*, LPS enhances mRNA levels in PeMPs both *in vitro* and *in vivo* (Figures 2 and 3).

**Figure 3.**
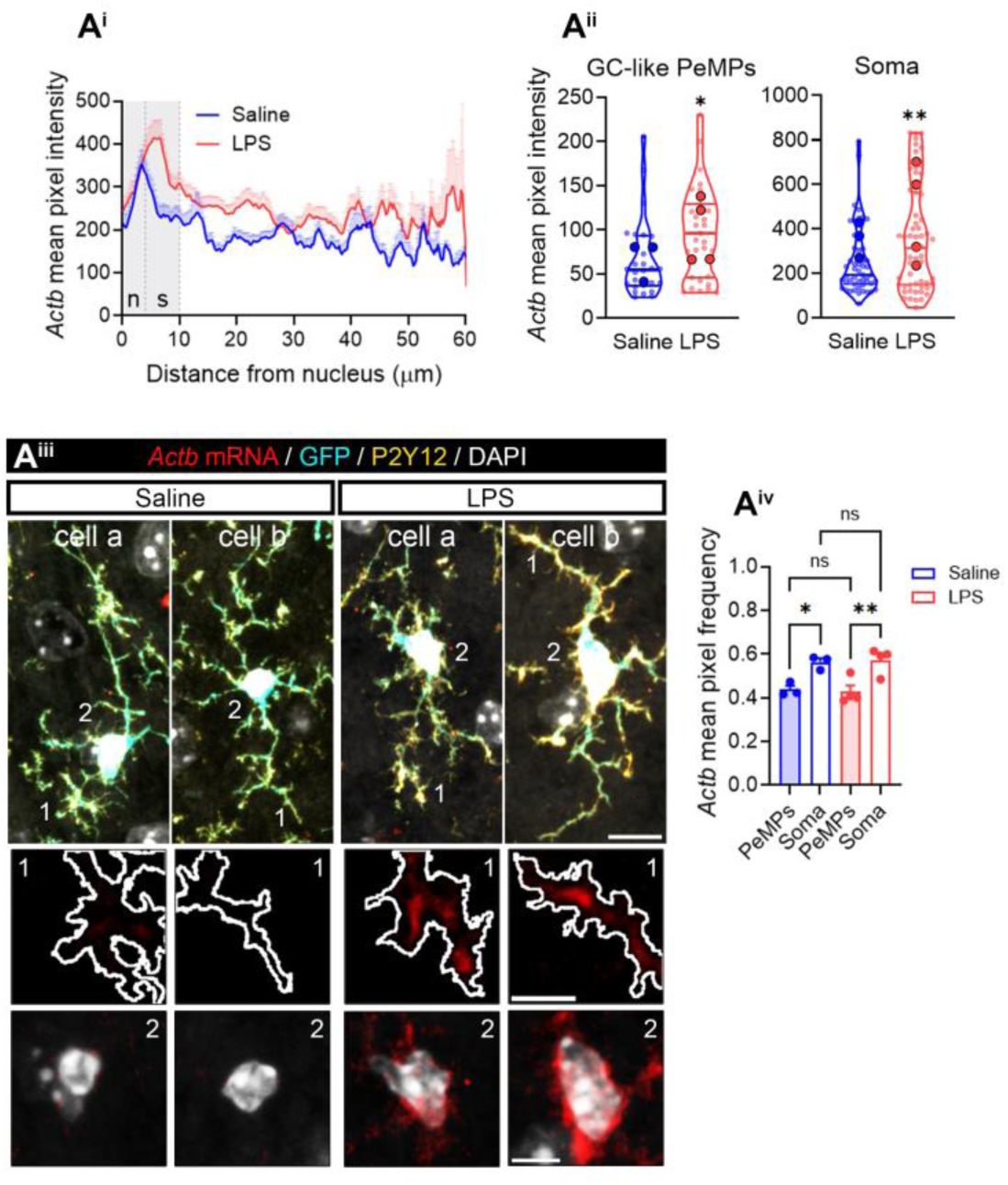
*Actb* levels are increased in LPS-injected mice. Linescans represent the distribution of the *Actb* FISH signal from the nucleus to the PeMPs in cortical microglia (positive for P2Y2 and GFP) from fms-EGFP 1-month old mice (A^i^). Violin plots represent the mean intensity of *Actb* in 26-31 sampled GC-like PeMPs (smaller dots) and 50-58 cell bodies (smaller dots) from 3-4 mice (larger dots). Two-tailed t tests. *p < 0.05; **p < 0.01 (A^ii^). Two cells per condition are shown as examples (cells a and b). (1) and (2) indicate GC-like PePMs and cell bodies represented in insets. Scale bars 10 µm (5 µm insets) (A^iii^). The bar graph shows the relative frequency distribution of *Actb* intensity in PeMPs and the soma in cortical microglia from 3-4 mice (n=3-4). One-way ANOVA followed by Holm-Šídák’s multiple comparison test. *p < 0.05; **p > 0.01 (A^iv^).

### LPS induces *Actb* and *Par3* localized translation in PeMPs

To address if *Actb* polarization upon acute LPS exposure *in vitro* was accompanied by changes in local translation we performed proximity ligation assays (PLA) combining specific antibodies against puromycin and β-actin protein. PLA enables the detection of newly synthesized target proteins during the timeframe of puromycin labelling ^35^. Since levels of newly produced proteins are typically lower in the processes than their soma-produced counterparts ^29^, to ensure the PLA signal was protein synthesis-specific we exposed cultured microglia to PBS or LPS for 30 minutes and co-incubated the cells with vehicle or with 40 µM of the protein synthesis inhibitor anisomycin. Nascent polypeptides were labelled with puromycin for the last 2 or 10 minutes of treatment. Levels of newly synthesized β-actin (Figures S4A and 4A) were blocked by anisomycin in LPS-but not in PBS-treated cells. For comparison, we also addressed *Par3* local translation. Newly synthesized Par3 results were similar to those obtained for newly synthesized β-actin (Figure S4B). Hence, we concluded that localized translation of both mRNAs of interest occurs in PeMPs in cells acutely treated with LPS. These results indicate that while *Actb* localization might be a limiting step for β-actin local synthesis in response to LPS, this inflammatory cue enhances local translation of pre-existing *Par3* mRNA in PeMPs, at least *in vitro*.

### RNA binding protein IMP1/ZBP1 drives *Actb* mRNA localization and localized translation in PeMPs

Zipcode binding protein 1 (IMP1/ZBP1) was the first RNA-binding protein (RBP) associated to the localization of *Actb* to fibroblast lamellipodia ^22^. Since then, IMP1-dependent asymmetric distribution of *Actb* has been described in many cell types, including neurons. Given the role of IMP1/ZBP1 in *Actb* mRNA localization we aimed to explore if it played a similar role in microglia.

First, we analyzed the expression of IMP1/ZBP1 in microglia and observed a granular pattern in the cytoplasm (Figure S4C^i^). A nonsignificant trend toward a decrease was found upon LPS exposure (Figure S4C^ii^) and no changes were observed in PeMPs (Figures 4Bi and 4B^ii^). However, IMP1/ZBP1-positive foci were restricted to the somatic region in control cells whereas upon LPS treatment IMP1/ZBP1 puncta showed more scattered distribution (Figures 4Biii and 4B^iv^), partially mirroring the change in *Actb* distribution upon LPS exposure *in vitro* (Figures 2Biv and 2B^v^). Conversely, IMP1/ZBP1 was generally increased in microglia *in vivo* in LPS-injected mice (Figures 4Ci and 4C^ii^), and despite these changes were not observed in GC-like PeMPs they did affect primary processes and the soma (Figure 4Ciii). Levels of this RBP were higher in the soma than in PeMPs in both saline- and LPS-injected animals (Figure 4Civ). These results again partially reproduced *Actb* mRNA pattern in response to systemic inflammation in mice (Figure 2C).

**Figure 4.**
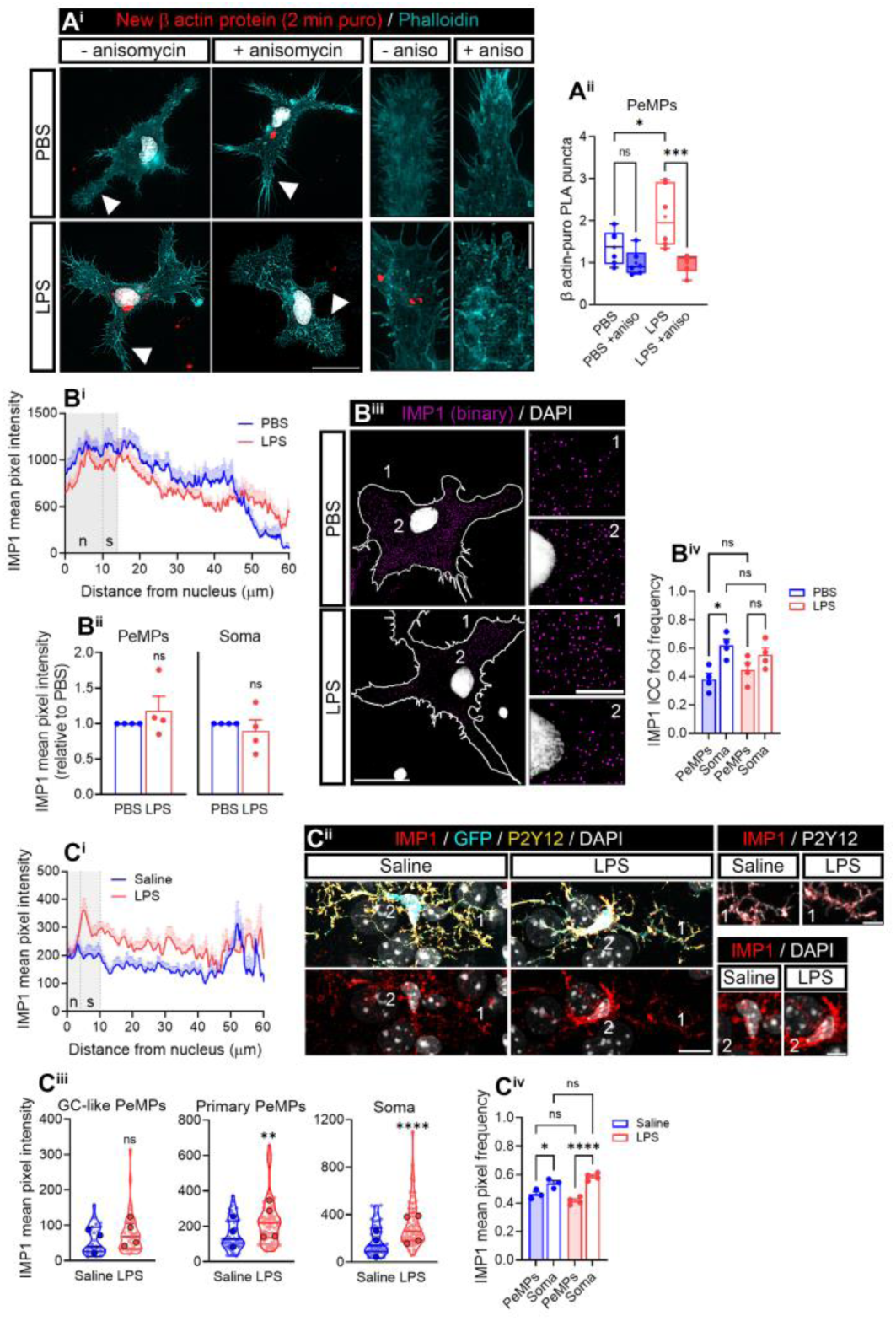
LPS enhances *Actb* localized translation and regulates RNA binding protein IMP1/ZBP1 in microglia. **(A)** New synthesis of β actin in microglial peripheral structures was assessed with a 2-minute puromycin pulse in PBS- and LPS-treated cells followed by proximity ligation assay (PLA) with antibodies against puromycin and β actin. As a negative control, cells were preincubated with the protein synthesis inhibitor anisomycin. Micrographs show the PLA signal y microglia labelled with phalloidin. Arrowheads indicate PeMPs depicted in insets (4 rightmost panels). Scale bars 20 µm (insets 10 µm) (A^i^). The box and whisker graph represents the average PLA puncta within lamellae in phalloidin-stained microglia treated with PBS or LPS in 6 independent cultures (n=6). One-way ANOVA followed by Holm-Šídák’s multiple comparison test. *p < 0.05; ***p < 0.001; ns, not significant (A^ii^). **(B)** IMP1/ZBP1 levels and relative distribution in microglia. Linescans represent the distribution of the IMP1 fluorescence signal from the nucleus to the PeMPs in 40 individual cells per condition (B^i^) Estimation plots show IMP1/ZBP1 levels in LPS-treated cells relative to controls in PeMPs and the soma measured in 4 independent experiments (n=4). Two-tailed t tests. n.s, not significant (B^ii^). Micrographs (binarized images) show the distribution of IMP1 in PBS- and LPS-treated cells. (1) and (2) indicate the PePMs and the perinuclear regions respectively represented in insets Scale bars, 20 µm (10 µm insets) (B^iii^). The box and whisker graph shows the relative frequency distribution of IMP1 foci in the periphery and the soma of microglia after treatment with PBS or LPS for 30 minutes in 4 independent experiments (n=4). One-way ANOVA followed by Holm-Šídák’s multiple comparison test. *p < 0.05; ns, not significant (B^iv^). **(C)** IMP1 levels measured by immunohistochemistry in cortical microglia (positive for P2Y2 and GFP) from fms-EGFP 1-month old mice injected with saline or LPS. Linescans represent the distribution of the fluorescent signal from the nucleus to the PeMPs in 55-72 individual cells per condition (C^i^). (1) and (2) indicate PePMs (both GC-like and primary processes) and cell bodies respectively shown in insets. Scale bars 10 µm (5 µm insets) (C^ii^). Violin plots represent the mean intensity of IMP1 in 28-36 sampled GC-like PeMPs (smaller dots), 55-72 primary PeMPs (smaller dots) and 55-72 cell bodies (smaller dots) from 3-4 mice (larger dots). Two-tailed t tests. **p < 0.01; ****p < 0.0001; n.s, not significant (C^iii^). The bar graph shows the relative frequency distribution of IMP1/ZBP1 intensity in PeMPs and the soma in cortical microglia from 3-4 mice (n=3-4). One-way ANOVA followed by Holm-Šídák’s multiple comparison test. *p < 0.05; ****p > 0.0001; n.s, not significant (C^iv^).

Since limiting the amount of IMP1/ZBP1 is sufficient to decrease *Actb* mRNA availability in axons ^24^, we sought to downregulate this protein by genetic silencing in microglia. We transfected cultured microglia with two nonoverlapping siRNAs against *Imp1* (siRNA #1 and siRNA #2) and used a non-targeting siRNA as a negative control. Whereas siRNA #1-transfected cells showed a trend towards a decrease in IMP1/ZBP1 levels after 24-hour transfection, siRNA #2 significantly downregulated the target protein both at 6 and 24 hours (Figure S4D). Thus, we further used siRNA #2, hereafter *Imp1* KD. *Actb* mRNA was significantly increased in PeMPs in response to LPS in control-transfected cells (ctrl KD, Figures 5Ai, left linescan, and 5A^ii^), in line with results obtained from untransfected cells (Figures 2Bi and 2B^ii^). Interestingly, the effect of LPS was blocked by *Imp1* KD (Figures 5Ai, right linescan, and 5A^ii^). Furthermore, *Actb* mRNA foci frequency was lower in the soma and higher in the periphery of LPS-compared to PBS-treated microglia in ctrl KD cells. *Imp1* KD led to a loss of *Actb* polarization toward PeMPs in LPS-treated microglia (Figures 5Bi-B^iii^).

Next, we sought to determine if *Imp1* KD also affected local β-actin synthesis in PeMPs. We performed PLA assays to detect newly synthesized β-actin following a 2-minute puromycin pulse. Like in untransfected cells (Figures S4A and 4A), acute LPS exposure led to an increase in local *Actb* translation in control-transfected cells, which was blocked by preincubation with 40 µM anisomycin. *Imp1* KD overall decreased β-actin PLA signal levels (two-way ANOVA p < 0.05) and no protein synthesis was detected neither in PBS-nor LPS-treated cells above levels observed in the presence of anisomycin (Figure 5C).

**Figure 5.**
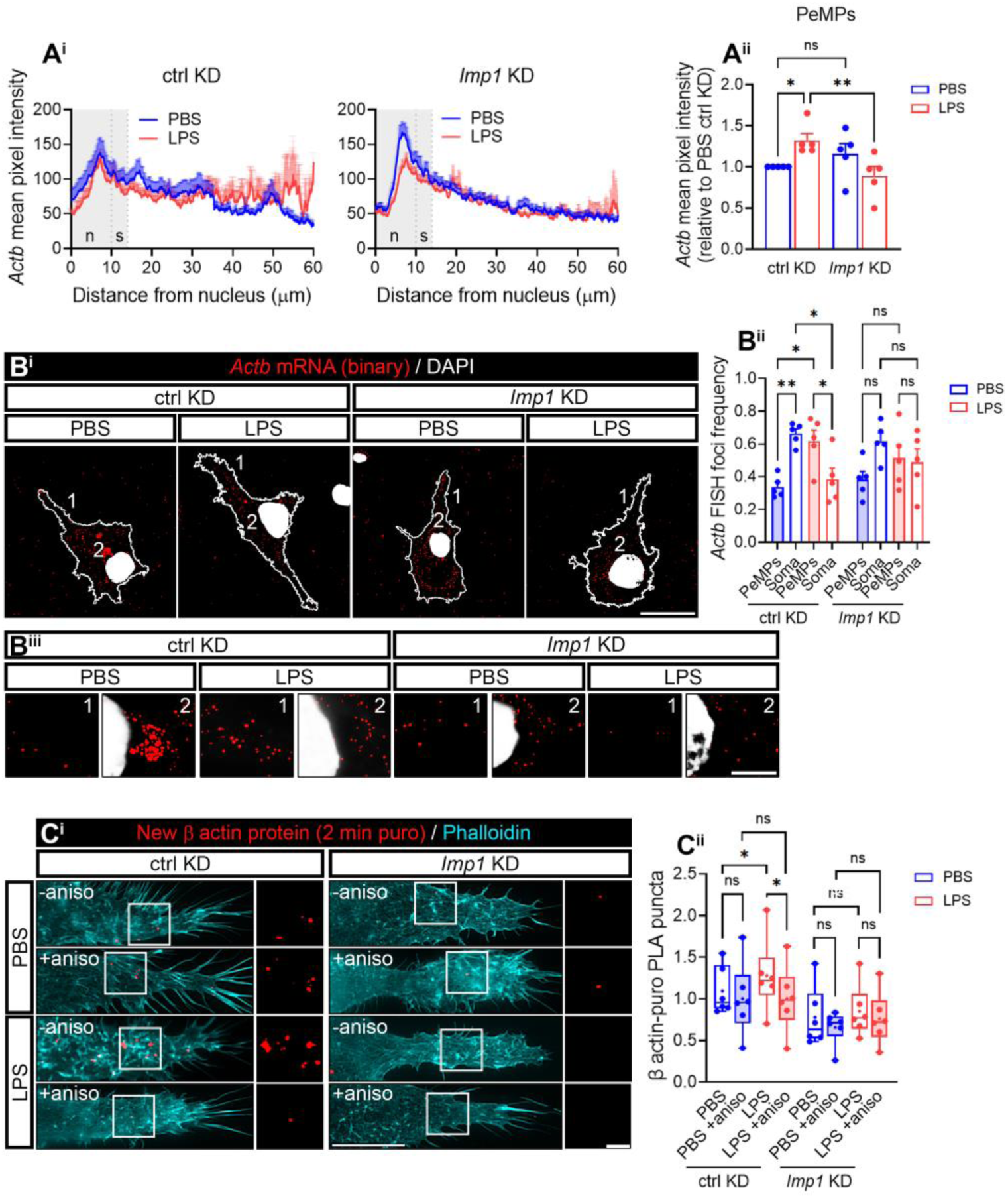
IMP1/ZBP1 regulates *Actb* mRNA localization and localized translation in microglia. **(A)** *Imp1* knockdown (KD) alters *Actb* mRNA localization towards the periphery of microglia. Linescans show the distribution of the *Actb* FISH signal from the nucleus to the PeMPs in 38-48 individual cells per condition (A^i^). The bar graph represents the relative *Actb* intensity (compared to control-transfected cells exposed to PBS) in PeMPs form microglia treated with PBS or LPS and transfected with a control (ctrl KD) or an *Imp1*-targeting *(Imp1* KD) siRNA in 5 independent experiments (n=5) One-way ANOVA followed by Holm-Šídák’s multiple comparison test. *p < 0.05; **p < 0.01; ns, not significant (A^ii^). **(B)** Analyses of the relative distribution of *Actb* foci in binarized images. (1) and (2) indicate the PePMs and the perinuclear regions respectively (B^i^) represented in insets (B^ii^). Scale bars, 20 µm (B^i^) and 10 µm (B^ii^). The relative frequency distribution of *Actb* foci in the periphery and the soma of microglia transfected with a control (ctrl KD) siRNA or an *Imp1* siRNA (*Imp1* KD) and exposed to PBS or LPS for 30 minutes in 5 independent experiments (n=5) is plotted in (B^iii^). Two-way ANOVA followed by Holm-Šídák’s multiple comparison test. *p < 0.05; **p < 0.01; ns, not significant. **(C)** *Imp1* KD blocks LPS-induced localized translation in microglia. Local β actin synthesis was assessed with a 2-minute puromycin pulse in PBS- and LPS-treated cells and transfected with control (ctrl KD) or *Imp1* siRNAs (*Imp1* KD), followed by PLA with antibodies against puromycin and β actin. As a negative control, cells were preincubated with the protein synthesis inhibitor anisomycin. Representative images of microglial PLA-labelled lamellae and stained with phalloidin are shown. Scale bar, 10 µm (insets, 5µm) (C^i^). The box and whisker graph represents the average of PLA puncta within lamellae in phalloidin-stained microglia cultured in 6 independent experiments (n=6) anayzed by two-way ANOVA followed by Holm-Šídák’s multiple comparison test. *p < 0.05; ns, not significant (C^ii^).

Altogether these results indicate that IMP1/ZBP1 regulates *Actb* mRNA localization to PeMPs and its localized translation in response to LPS.

### LPS induces morphological changes in microglia and enhances random motility, and polarized PeMP extension in an IMP1/ZBP1-dependent manner

*Actb* mRNA is the prototypical example of a localized transcript in many eukaryotic cells and its local translation is involved in cytoskeletal rearrangements, cell polarity and lamellar migration towards attractant cues ^36^. In our acute inflammation model, IMP1/ZBP1 downregulation alters *Actb* mRNA localization to PeMPs and its localized translation (Figure 5) *in vitro*. Thus, our next step was to address any morphological changes in microglia occurring in response to LPS treatment and when local β actin availability was limited in microglial processes by *Imp1* knockdown. To that end we first performed morphological analyses in cortical microglia from fms-EGFP mice injected with saline or LPS. Upon LPS administration, microglia acquired a bushier morphology with fine processes emerging from primary processes close to the soma (Figure 6Ai, upper panels). These results are consistent with a higher morphological complexity as quantified by 2D Sholl analyses (Figure 6Aii, upper graph) and with the increase in mean intersections observed in LPS-compared to saline-injected mice (Figure 6Aiv). However, we noticed no changes in PeMP extension measured as the maximum cell radius (Figure 6Aiii). Interestingly, morphological changes driven by LPS were no longer observed in mice that also received anisomycin (Figure 6Ai, lower panels. Figure 6Aii, lower graph), and mean intersections were significantly decreased in LPS-injected animals administered with this drug (Figure 6Aiv). These results suggest that LPS-induced morphological complexity requires protein synthesis. To determine whether IMP1/ZBP1-dependent *Actb* local translation could be responsible for microglial morphological changes in response inflammation, we transfected primary cultures with a control or with an *Imp1*-targeting siRNA. LPS did not alter PeMP length or width neither in untransfected cells (data not shown) nor in transfected microglia (Figure S5A), however it did modify filopodia distribution along the lamellar surface. We observed that most of fine processes were localized at the base of lamellae in PBS-treated microglia and redistributed more homogenously upon LPS exposure when cells were transfected with a control siRNA for 24 hours. Conversely, filopodia distribution was similar in both experimental conditions in *Imp1*-transfected cells. These results suggest that acute LPS treatment leads to the rapid rearrangement of filopodia in an IMP1/ZBP1-dependent manner (Figure 6B). We then analyzed PeMP motility by life cell imaging in cells treated with vehicle or LPS for 0, 5, 10, 15, 20 and 30 minutes. Cell visualization was performed for 2 minutes (5-sec cycles) at each time point. LPS enhanced PeMP motility after a 10-minute treatment (Figures 6Ci and 6C^ii^), however no net protrusion was observed (Figure S5C). To determine if IMP1/ZBP1 is required for LPS-induced motility, we transfected cells with siRNAs and, again, performed live imaging. In line with results from untransfected cells, LPS also increased random (undirected) PeMP motility in control-transfected cells and this effect was blocked by *Imp1* KD (Figures 6Ciii, 6C^iv^ and S5D).

**Figure 6.**
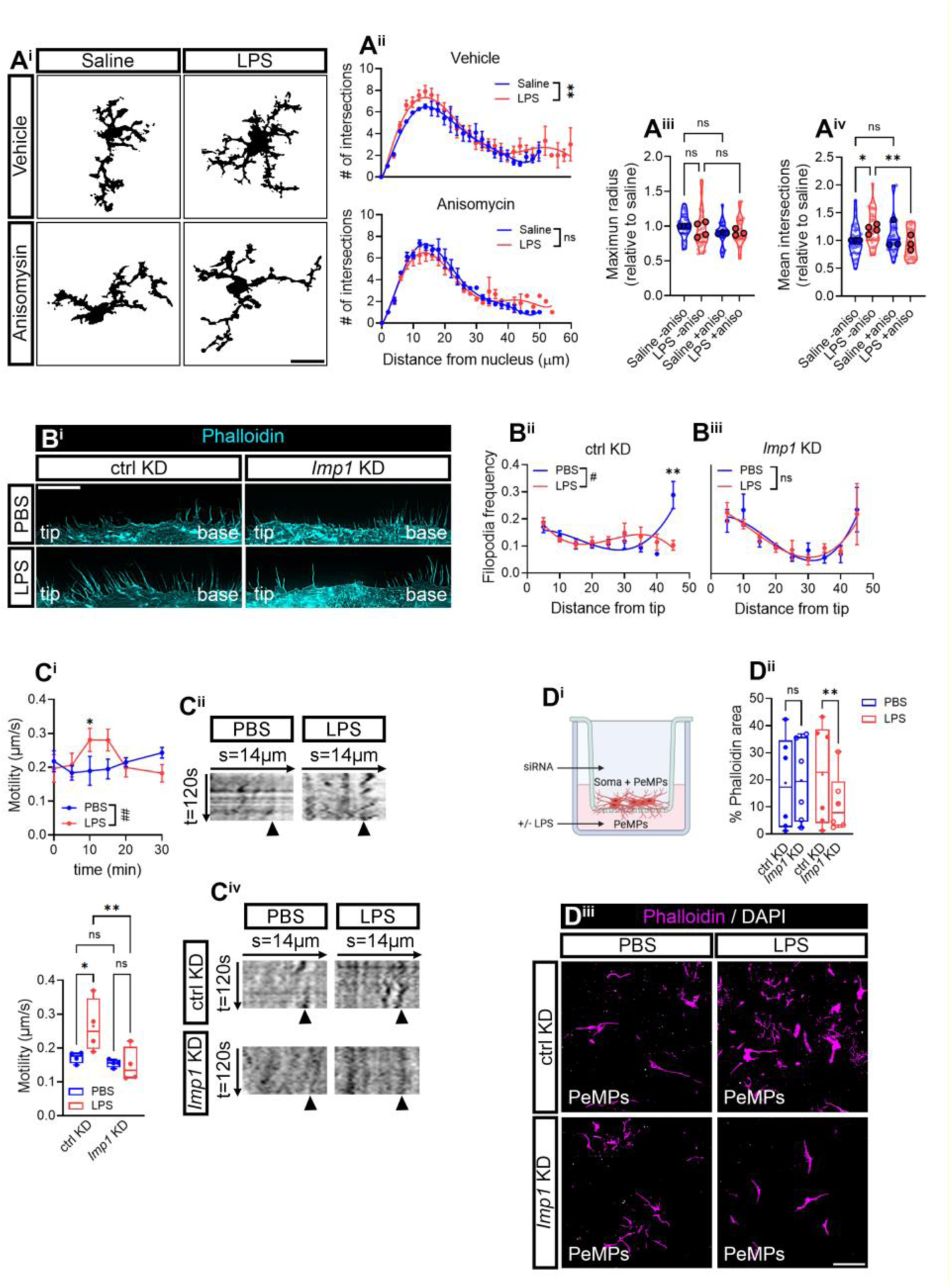
LPS-induced PeMPs morphological changes depend on IMP1/ZBP1 levels. **(A)** 2D Sholl analyses of cortical microglia from fms-EGFP 1-month old mice injected with saline or LPS and co-injected with vehicle or anisomycin. Micrographs exemplify binarized cells used for Sholl analyses. Scale bar 20 µm (A^i^). Graphs represent Sholl analyses performed in 3-4 animals per experimental group (n=3-4). For simplification purposes animals injected with vehicle (upper graph) or anisomycin (lower graph) are represented separately, although statistical analysis using two-way ANOVA with Holm-Šídák’s *post hoc* test was used taking all experimental groups into account. **p < 0.01; n.s, not significant (A^ii^). PeMP extension in response to LPS *in vivo* was measured as the cell maximum radius of 25-36 cortical microglia (smaller dots) from 3-4 animals (larger dots). One-way ANOVA followed by Holm-Šídák’s multiple comparison test. ns, not significant (A^iii^). Mean intersections from 25-36 cortical microglia (smaller dots) measured in 3-4 animals (larger dots) are represented in violin plots in (A^iv^). One-way ANOVA followed by Holm-Šídák’s multiple comparison test. *p < 0.05; **p <0.01; n.s, not significant. **(B)** Distribution of filopodia along microglial lamellae in control- (ctrl KD) and *Imp1* siRNA-transfected cells (*Imp1* KD) exposed to PBS or LPS for 30 mins. Representative micrographs depicting filopodia distribution in phalloidin-stained microglia are shown. Scale bar, 10 µm (B^i^). Graphs represent the relative filopodia distribution of microglia cultured in 6 independent experiments (n=6). Two-way ANOVA followed by Holm-Šídák’s multiple comparison test. #p < 0.05 (factor interaction); *p < 0.05; ns, not significant. (B^ii^). **(C)** Random PeMP motility measured in response to LPS in untransfected- and siRNA-transfected cells. Live cell imaging was performed in microglia exposed to PBS or LPS for different times and PeMP motility was assessed for 2 minutes every 5 seconds. Time-course graphs represent PeMP motility in response to PBS or LPS in 4-6 independent cultures from untransfected cells (n=4-6). Two-way ANOVA followed by Holm-Šídák’s multiple comparison test. #p < 0.05 (factor interaction); *p < 0.05; n.s, not significant (C^i^). Kymographs show the representative oscillations of microglial lamellae during 2 min imaging (5 sec cycles) after a 10-minute treatment with vehicle or LPS (C^ii^). Box and whisker graph show the mean PeMP motility in control-transfected (ctrl KD) and *Imp1* KD cells exposed to vehicle or LPS for 10 minutes from 4 independent cultures (n=4). One-way ANOVA with Holm-Šídák’s *post hoc* test (C^iii^). Kymographs show the representative oscillations of microglial lamellae during 2 min imaging (5 sec cycles) after a 10-minute treatment with PBS or LPS in control- or *Imp1*-siRNA transfected microglia (C^iv^). **(D)** To address lamellar migration, we restricted the soma to the upper compartment by culturing cells in 1-µm-diameter transwell membranes. PBS or LPS were applied to the bottom to the lower compartment for 30 minutes (D^i^). The bar graph represents the relative coverage of phalloidin staining at the lower side of the membrane. Analyses were performed in 6 independent experiments (n=6) by one-way ANOVA followed by Holm-Šídák’s multiple comparison test. **p < 0.01; ns, not significant; test (D^ii^). Representative images of lamellae migrated to the lower side of the membrane are shown. Scale bar, 50 µm (D^iii^).

All experiments reported in this section thus far were performed either by systemic injection *in vivo* or bath application of vehicle or LPS to cells *in vitro*. In this context PeMP motility is unlikely directed as treatments are not focal, which could explain why we were not able to observe PeMP extension Thus, as a next step we attempted to focalize the treatments by culturing cells in transwells. Cells were seeded on top of a 1 µm-pore membrane and LPS or vehicle were applied to the lower compartment (Figure 6Di). As mentioned before, this culture setup enables microglial PeMP extension toward the lower compartment restricting cell body migration. In this case, LPS did enhance PeMP polarized migration measured as the area covered by F-actin in untransfected and control-transfected cells (Figure S7A) and this effect was significantly reduced upon IMP1/ZBP1 downregulation in LPS-treated microglia (Figures 6Dii and 6D^iii^). Interestingly, if cell bodies were allowed to move toward the lower compartment by culturing microglia on 3 µm-pore membranes, neither LPS nor IMP1/ZBP1 affected cell migration (Figures S7B and S7C), suggesting that PeMP motility and cell body migration might be regulated by different mechanisms, being the latter LPS-and IMP1-independent.

### IMP1/ZBP1 and *Actb* mRNA are enriched in phagocytic pouches, and IMP1 downregulation impairs microglial phagocytosis

Phagocytosis is one of the main functions of microglia to restore brain homeostasis by removing cell debris and damaged cells upon inflammation, and it is driven by PeMPs *in vivo*. During phagocytosis, PeMPs rearrange their cytoskeleton and form specialized peripheral structures termed phagocytic pouches ^27^. Interestingly, Vasek and colleagues have recently reported enrichment of transcripts important for phagocytosis in a PeMP localized translatome, and that protein synthesis is required for efficient microglial phagocytosis ^12^. Given that local β-actin production is required for rapid cytoskeletal remodelling in response to external guidance cues ^20^, and that our own data indicate that *Imp1* KD decreases the levels of *Actb* in PeMPs (Figure 5), filopodia distribution (Figure 6B), PeMP motility (Figure 6C) and LPS-dependent polarized PeMP extension (Figure 6D), we sought to analyze the relevance of IMP1/ZBP1 on phagocytosis.

As a first step we analyzed the presence of phagocytic pouches *in vivo* to determine if active ribosomal protein Rsp6, IMP1/ZBP1 and *Actb* mRNA were regulated in these peripheral structures in response to LPS. We focused on the subgranular zone of the dentate gyrus where naturally occurring cell death associated to neurogenesis can be detected in 1-month-old mice ^37^. Indeed, apoptotic cells surrounded by microglial pouches were readily present in this brain region (p in micrographs shown in Figures 7A and 7B). Thus, we quantified levels of pS6 in phagocytic microglia and despite no changes were observed in LPS-compared to saline-injected mice (data not shown), a trend towards an increase in pouches compared to the soma was detected in the latter, while this increase was significant in animals that received LPS (Figure 7A). These results suggest an enrichment of active protein synthesis in phagocytic PeMPs. Importantly, both IMP1/ZBP1 and *Actb* mRNA showed and overlapping pattern in phagocytic microglia (Figure 7B), being both selectively increased in pouches (Figure 7Ci and 7C^iii^) but not in the soma (see linescans Figures 7B) in response to LPS. Indeed, both markers were enriched in pouches compared to the soma in LPS-injected animals (Figures 7Cii and 7C^iv^) and a significant positive correlation among them was observed (Figure 7Cv). These results could indicate that IMP1/ZBP1 drives the localization of *Actb* mRNA to phagocytic PeMPs *in vivo*, especially in a neuroinflammatory environment. These data were consistent with a significant increase in pouch diameter observed in LPS-injected mice exposed to the protein synthesis inhibitor anisomycin (Figure 7D). With these results it is tempting to speculate that IMP1-dependent local protein synthesis, including that of β-actin, is required for efficient phagocytosis by microglia.

**Figure 7.**
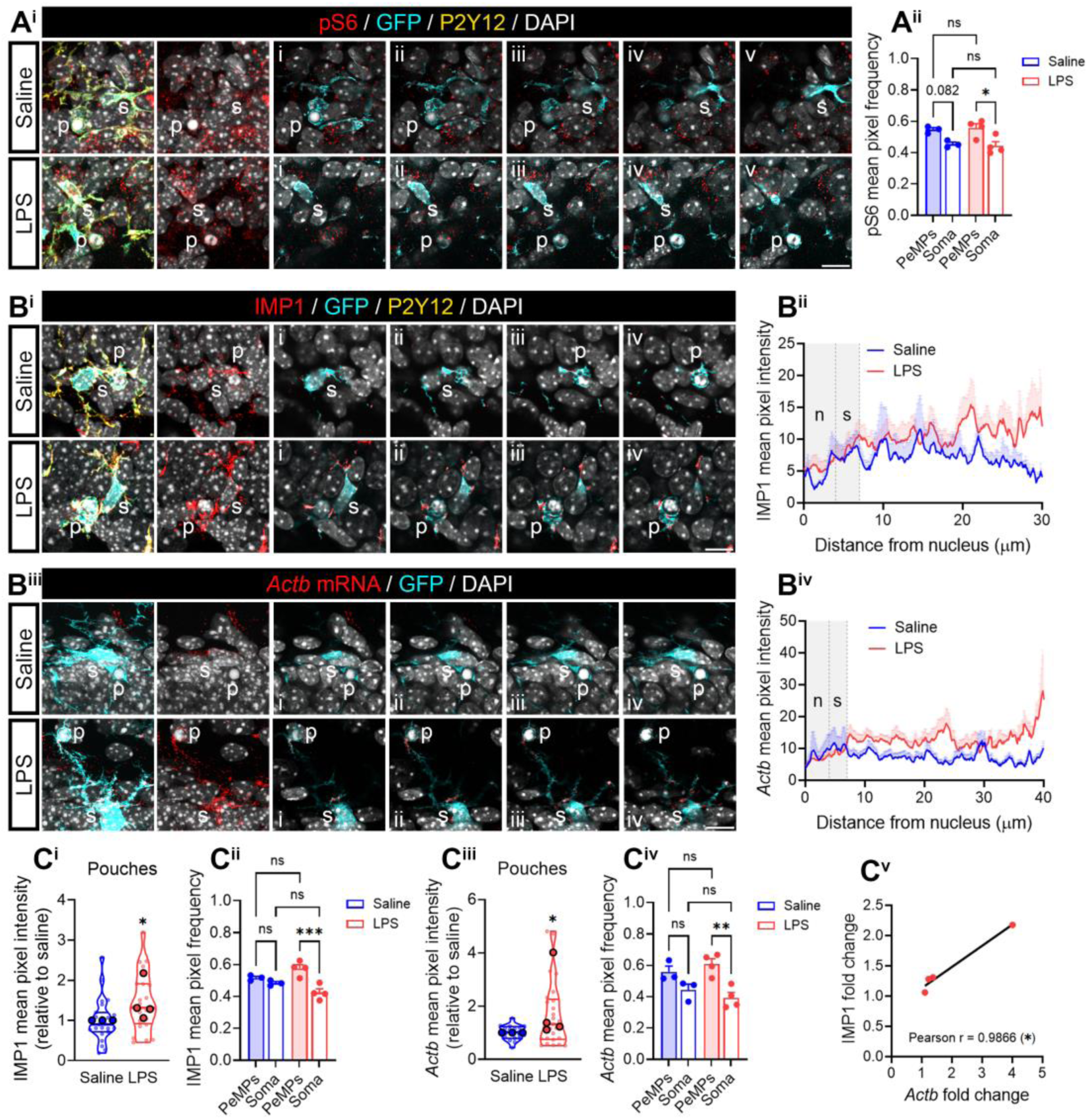
LPS induces the enrichment of pS6, IMP1 RBP and *Actb* mRNA in phagocytic pouches. **(A)** S6 phosphorylation levels in phagocytic microglia in the dentate gyrus. Representative micrographs show pS6 immunostaining in the cell body (s) and in phagocytic PeMPs (p) of microglia from saline- and LPS-injected mice. A series of 5 consecutive optical sections (i-v) are shown. Scale bar 10 µm (A^i^). The graph shows the relative frequency distribution pS6 intensity in phagocytic PeMPs and the soma in hippocampal microglia from 3-4 mice (n=3-4). One-way ANOVA followed by Holm-Šídák’s multiple comparison test. *p < 0.05; n.s, not significant (A^ii^). **(B)** IMP1 and *Actb* mRNA levels in phagocytic microglia in the dentate gyrus. Representative micrographs of IMP1/ZBP1 immunostaining in the cell body (s) and in phagocytic PeMPs (p) of microglia from saline- and LPS-injected mice. A series of 5 consecutive optical sections (i-v) are shown. Scale bar 10 µm (B^i^). Linescans represent IMP1 fluorescent signal from the nucleus to phagocytic pouches in 8-12 individual cells per condition (B^ii^). Representative micrographs of *Actb* FISH signal in the cell body (s) and in phagocytic PeMPs (p) of microglia from saline- and LPS-injected mice are shown. 5 consecutive optical sections are represented (i-v). Scale bar 10 µm (B^iii^). Linescans depict *Actb* FISH fluorescent signal from the nucleus to phagocytic pouches in 12-18 individual cells per condition (B^iv^). **(C)** LPS injection increases IMP1 and *Actb* mRNA levels in phagocytic pouches *in vivo*. Violin plots summarize the relative changes in IMP1/ZBP1 levels (LPS vs saline) observed in phagocytic PeMPs from 23-24 cells (smaller dots) measured in the hippocampus of 3-4 mice (larger dots) (C^i^). The graph shows the relative frequency distribution of IMP1/ZBP1 intensity in pouches and the soma in l microglia from 3-4 mice (n=3-4). One-way ANOVA followed by Holm-Šídák’s multiple comparison test. ***p > 0.001; n.s, not significant (C^ii^). Violin plots summarize the relative changes in *Actb* mRNA levels (LPS vs saline) observed in phagocytic PeMPs from 16-25 cells (smaller dots) measured in the hippocampus of 3-4 mice (larger dots) (C^iii^). The bar graph shows the relative frequency distribution of *Actb* intensity in pouches and the soma of microglia from 3-4 mice (n=3-4). One-way ANOVA followed by Holm-Šídák’s multiple comparison test. **p > 0.01; n.s, not significant (C^iv^). The correlation between the fold change (LPS vs saline) in IMP1 and *Actb* levels in pouches is quantified in (C^i^).

Thus, to determine to what extent IMP1/ZBP1 is actively involved in phagocytosis, we treated cultured microglia with PBS or LPS for 30 minutes prior to adding apoptotic cells. We fed cells with human SH-SY5Y vampire red transgenic apoptotic neurons for 1 hour and analyzed the percentage of microglia with vampire-positive particles inside. Around 60% of untransfected microglia (Figure S8A^i^) and 70% of control-transfected microglia had engulfed apoptotic neurons both in control and LPS-induced conditions (Figure 8Di). Additionally, *Imp1* KD had no effect on the proportion of phagocytic microglia (Figure 8Ai). However, particles engulfed by cells exposed to LPS were significantly smaller (Figure S8). *Imp1* KD increased particle size (Figures 8Aii and 8A^iii^) and altered the number of particles per cell in LPS-treated microglia, reducing the percentage of microglia with a higher number of engulfed particles, which were usually smaller (8B), suggesting that despite IMP1/ZBP1 is unlikely involved in engulfment it might play a role in degradation of phagocytosed cells.

**Figure 8.**
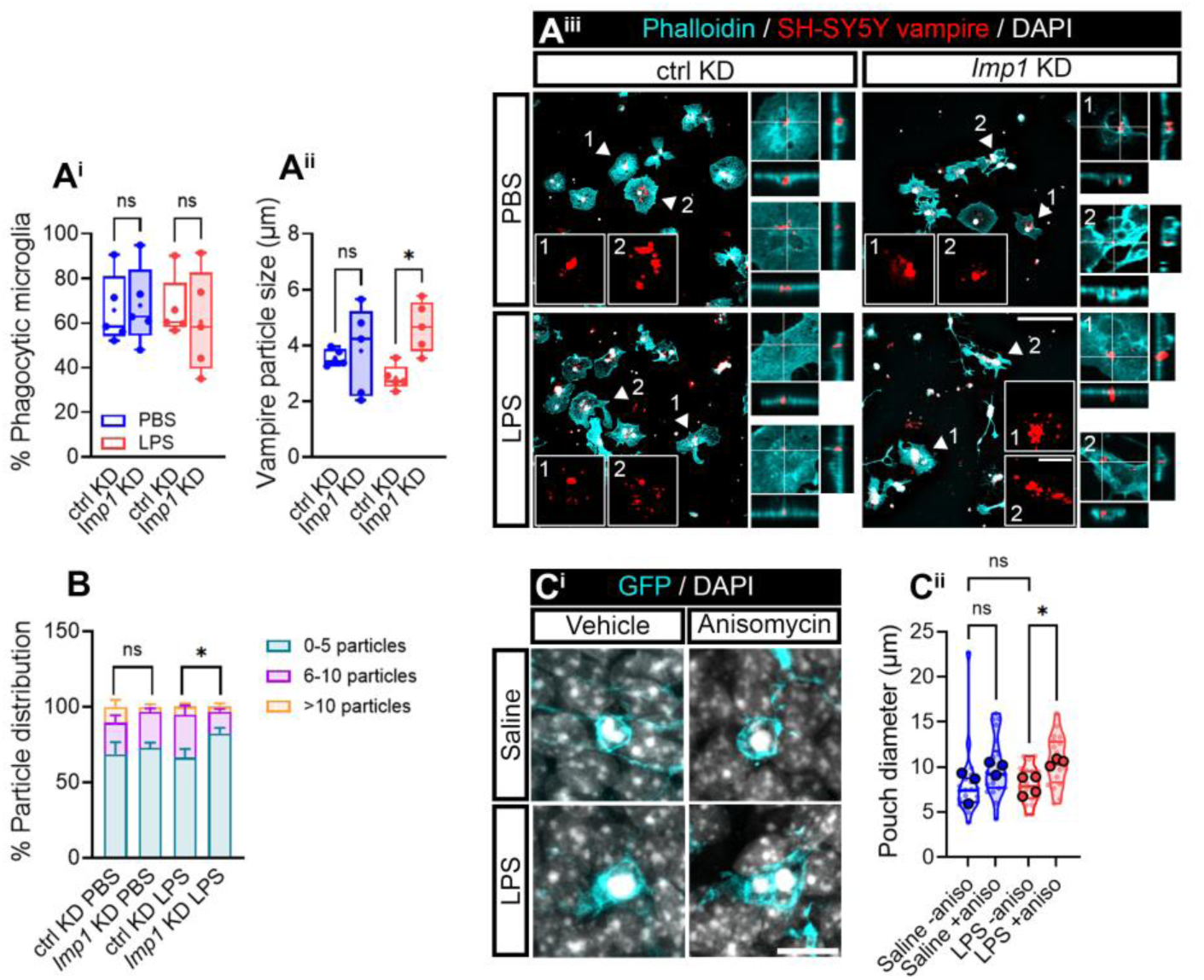
*Imp1* KD increases the size of phagocytosed particles. **(A)** Percentage of control- (ctrl KD) and *Imp1*-transfected (*Imp1* KD) phagocytic microglia after treatment with SH-SY5Y apoptotic neurons labelled with vampire for 1 hour after by exposure to PBS or LPS for 30 minutes in control-transfected and *Imp1* KD cells (A^i^) and vampire particle size in following the same experimental setup (A^ii^). Both graphs correspond to 5 independent experiments (n=5). Two-way ANOVA followed by Holm-Šídák’s multiple comparison test was performed. *p < 0.05; ns, not significant. Representative micrographs of vampire-positive neurons (red) inside phalloidin-stained microglia are shown. Insets depict examples of vampire particles inside phagocytic microglia. Zoomed orthogonal views are also represented for each experimental condition. Scale bar, 100 µm; 10 µm in insets and orthogonal view (A^iii^). **(B)** Frequency distribution of phagocytic microglia with 1-5, 6-10 and >10 apoptotic neurons inside, following transfection with control (ctrl KD) and *Imp1* (*Imp1* KD) siRNAs and treated with PBS or LPS for 30 minutes. The graph represents the mean of 5 independent experiments (n=5). Two-way ANOVA followed by Holm Sidak’s multiple comparison test was performed, *p < 0.05; n.s, not significant. **(C)** Phagocitic pouches are enlarged in mice treated with LPS and exposed to the protein synthesis inhibitor anisomycin. Representative micrographs of GFP-positive pouches with engulfed apoptotic cells are shown. Scale bar 10 µm (C^i^). Violin plots summarize the pouch diameter of 20-24 phagocytic PeMPs (smaller dots) measured in the hippocampus of 3-4 mice (larger dots) (C^ii^).

In summary, our data confirm that microglia are able to produce proteins locally at their PeMPs ^12^. In addition, we identified *Actb* as one of the transcripts whose PeMP localization and translation is induced by the inflammatory cue LPS. As in other polarized cells, *Actb* localization to microglial PeMPs is regulated by the RBP IMP1/ZBP1, and our results suggest that IMP1/ZBP1-dependent *Actb* localization to PeMPs and its local translation is involved in fundamental microglial functions, including filopodia rearrangements, PeMPs motility and polarized migration, and phagocytosis.

## DISCUSSION

Here we report that LPS triggers within minutes the translation in PeMPs of two transcripts involved in cell polarization, cytoskeletal rearrangement and migration, one of them being *Actb*. Downregulation of the RBP IMP1/ZBP1 impairs *Actb* localized translation and leads to defects in LPS-dependent filopodia rearrangements, PeMP migration and phagocytosis. Thus, the microglial response to acute inflammation functionally requires local protein synthesis within PeMPs.

This work was motivated by evidence that microglia, like many other highly polarized cells, sense local changes in their environment with their peripheral processes which respond independently from their cell soma. Local PeMP responses, including process migration or phagocytosis are required to maintain brain homeostasis, which might be imbalanced by pathological insults like neuroinflammation ^17,27^. Thus, microglia rely on efficient signalling initiated at PeMPs ^17^, which we reasoned could be accompanied by changes in the local proteome. Local proteomes can be shaped by proteins transported from the soma or by those synthesized locally in PeMPs. Indeed, in neurons, oligodendrocytes, astrocytes and radial glia, local protein synthesis is required to maintain local protein homeostasis and for cells to accurately respond to environmental cues, as proteins are produced only when and where they are needed ^3,10^. In the case of microglia however, local translation had not been described until recently ^12^, and, to our knowledge this is the first report that describes local protein synthesis in PeMPs in response to inflammation. Given the limited information from microglial cells, we sought to explore this molecular mechanism based on the similarities between axonal growth cones and PeMPs.

PeMPs highly resemble axonal growth cones in terms of their morphology and behaviour. Both contain fine processes, termed filopodia, which survey their environment. Upon environmental changes that induce migratory responses, filopodia extend or collapse and drive large processes (lamellipodia) towards chemoattractants or away from chemorepellents. During development growth cones respond to NGF and netrin-1 by locally translating *Par3*, which leads to axonal outgrowth ^33^. Similarly, β-actin is locally produced in response to netrin-1 and enables actin cytoskeleton polymerization and reorganization leading to axon elongation ^20^. Hence, we asked if *Actb* and *Par3* mRNAs where also locally translated in PeMPs and to what extent were microglia to inflammation. Our results indeed confirmed that both transcripts polarized toward PeMPs and were locally translated upon inducing an acute inflammatory state in microglia. Interestingly, neither *Actb* nor *Par3* could be detected in PeMPs when challenging microglia with LPS for 24 hours, although we could not rule out *Par3* translation at this time point. These results suggest that translation of a distinct cohort of localized mRNAs might be required for microglia to differentially adapt to acute and chronic insults. This is reminiscent of the axonal response to pathological stimuli. For instance, long exposure of axons to Aβ oligomers, involved in Alzheimeŕs disease, drives the local translation of the transcription factor *Atf4*, which is not triggered by acute Aβ treatment. Importantly, late axonal synthesis of ATF4 itself depends on an early local production of vimentin ^30^. Thus, distinct waves of local translation are required for cells to adapt to the nature and duration of stimuli, and local transcriptomes and translatomes likely need to be dynamically regulated in time to continuously support the accurate response of peripheral processes, including PeMPs, to environmental changes.

For local translation to occur, mRNAs and components of the protein synthesis machinery must be transported to the target peripheral compartment. Following transcription, RBPs bind mRNAs through the recognition of cis-localization elements typically present in the 3’ untranslated region (3’UTR) of the transcripts ^38^. The resulting ribonucleoprotein complexes (RNPs) are transported by molecular motors in a translationally repressed state. Once localized to their destination and upon stimulation mRNAs are released from the RNPs, associate to the localized protein synthesis machinery and translate into proteins ^5^. In order to address the functional relevance of local translation in microglia, we downregulated IMP1/ZBP1, which was the first RPB associated to the localization of *Actb* to fibroblast lamellipodia ^22^. Since then, IMP1-dependent asymmetric distribution of *Actb* and its involvement in local translation have been described in many cell types, including neurons. Importantly, *Imp1* haploinsufficiency decreases *Actb* mRNA localization to axons and limits the regeneration of peripheral nerves upon injury ^24^. Thus, we reasoned that limiting the availability of IMP1/ZBP1 in microglia would affect LPS-dependent β-actin synthesis in PeMPs and it could uncover a functional relevance of local translation in the microglial response to inflammation. Indeed, genetic silencing experiments did confirm that IMP1/ZBP1 is required for *Actb* mRNA polarization to and localized translation within PeMPs in response to LPS. More importantly, acute LPS exposure seemed to induce changes only in the behaviour of PeMPs. For instance, LPS drove filopodia rearrangements along lamellipodia but did not affect lamellar morphology. On the other hand, whole cell migration was not affected by LPS treatment, which conversely induced PeMP directed motility when restricting somatic migration. It is precisely in both these contexts that IMP1/ZBP1-dependent local β-actin synthesis seemed to play a relevant role, evidencing the dependency of microglia on local translation to rapidly react to environmental changes sensed by their peripheral processes. One very relevant function carried out by microglia is phagocytosis, which is essential to restore brain homeostasis by removing cell debris and damaged cells upon inflammation. Phagocytosis is performed by PeMPs *in vivo* and requires rapid morphological changes of processes to form the so called phagocytic pouches, independently from the cell soma ^27^. Importantly, ribosome-bound transcripts involved in phagocytosis are enriched in PeMPs *in vivo* and *de novo* protein synthesis is required for the formation of phagocytic pouches *ex vivo* ^12^. In line with these observations our results showed IMP1/ZBP1 downregulation impaired LPS-dependent phagocytosis of apoptotic neurons, confirming the functional relevance of RBP-mediated local protein synthesis in response to inflammation.^10^. Importantly, many genetic neurological disorders, such as fragile X syndrome (FXS), amyotrophic lateral sclerosis (ALS), spinal muscular atrophy (SMA) or Hungtintońs disease are characterized by mutations in RBPs, and defects in RNA localization and/or local translation are common to all of them. IMP1/ZBP1 has been reported to bind several mRNAs encoded by autism spectrum disorder-genes ^39^ and our findings indicate that limiting IMP1/ZBP1 availability disrupts mRNA localization and translation in MPPs, and leads to dysfunctional microglia. Thus, it is tempting to speculate that defects in RBPs not only impair local protein synthesis in neurons, but also in microglia contributing to neurological disorders.

One of the limitations of this study is that mechanistic experiments have been performed *in vitro* using siRNAs as the only genetic tools for local translation inhibition, one of the reasons being the difficulty to genetically-manipulate microglia by transfection and transduction both *in vitro* and *in vivo* ^40^. Now that microglia are finally in the arena of the local translation field, it will be interesting to develop novel tools to address to what extent defects in RBPs and dysregulated local protein synthesis lead to microglia dysfunction and contribute to the development of brain pathologies in disease models. However, our proof-of-concept work already provides an unprecedented mechanistic insight into local translation regulation in microglia and points towards deficient response to inflammation when local β-actin synthesis in PeMPs is impaired.

## MATERIALS AND METHODS

### Animals

All animal protocols followed the European directive 2010/63/EU and were approved by the UPV/EHU ethics committee. For microglial cultures, Sprague Dawley rats were bred in local facilities and brains were obtained from P0-P2 postnatal rats. All *in vivo* experiments were performed in 1-month old mice expressing the enhanced green fluorescent protein under the control of the colony stimulating factor 1 receptor (fms-EGFP) (MacGreen, B6. Cg-Tg (Csf1r-EGFP) 1 Hume/J; Jackson Laboratory stock #018549). The fms-EGFP mouse colony, which expresses GFP at the same physiological levels as CSF1R was established in Achucarro by Dr. Amanda Sierra.

### Primary microglia cultures

Cortical microglial cells were isolated from mixed glial cultures. Briefly, brain hemispheres were dissected from postnatal Sprague Dawley rats (P0-P2) and dissociated with 0.25% trypsin (Sigma Aldrich, Merck, Darmstadt, Germany) and 0.004% DNAse (Sigma Aldrich) for 15 minutes at 37°C. Trypsinization was stopped by adding glial plating medium containing IMDM (Gibco, Thermo Fisher Scientific, Waltham MA, USA), 10% fetal bovine serum Hyclone (Cytiva, Thermo Fisher Scientific) and a mixture of antibiotics and antimycotics (Gibco). Cells were homogenized and centrifuged at room temperature for 6 minutes at 1200 rpm, resuspended in 1 ml plating medium and mechanically dissociated using 21G and 23G needles. Cells were centrifuged at room temperature for 6 minutes at 1200 rpm, resuspended in plating medium and seeded onto 75 cm^2^ flasks (BioLite, Thermo Fisher Scientific). Cultures were maintained at 37 °C in a 5% CO^2^ humidified incubator. After 1 day in vitro (DIV) the medium was replaced with glucose Dulbecco’s Modified Eagle’s Medium (DMEM, Gibco) containing 10% fetal bovine serum (Sigma Aldrich), 1 U/ml penicillin, 1 µg/ml streptomycin and 292 µg/ml glutamine (all from Gibco). The medium was replaced every 3 days.

For microglia isolation, 10-17 DIV mixed glial culture flasks were agitated at 180 rpm for 4h at 37 °C. The medium was collected and centrifuged at room temperature for 6 minutes at 1200 rpm. Cells were resuspended in serum-free medium containing Neurobasal, 1X B-27 supplement and 2 mM L-glutamine (all from Gibco). Microglia were seeded on Poly-D-lysine (PDL)-coated coverslips (40.000-80.000 cells/cm^2^), on 1-and 3-µm pore transwells (60.000-80.000 cells/cm^2^) or in 35 mm µ-Dish polymer coverslip bottom dishes (Ibidi, Gräfelfing, Germany) (90.000-100.000 cells/cm^2^). Treatments were performed at 12-14 DIV or 19-21DIV.

### *In vitro* pharmacological treatments

Lipopolysaccharides from *Escherichia coli* (LPS, BioXtra, Sigma Aldrich) were added to cells at 150 ng/ml for 30 minutes or 24 hours. PBS was used as vehicle control.

For puromycilation assays cells were exposed to 2 µM puromycin dihydrochloride from *Streptomyces alboniger* for 2, 10 or 30 minutes (Sigma Aldrich). Water was used as a vehicle control. Whenever stated, cells were preincubated for 30 minutes with anisomycin (Sigma Aldrich) at 40 μM to block protein synthesis.

### *In vivo* pharmacological treatments

1-month old MacGreen mice (both male and female) were intraperitoneally administrated with LPS with (1 mg per kg of body mass. BioXtra) or saline solution (NaCl 0.9%) daily for 4 consecutive days. Another group of animals received intraperitoneal injections of saline (NaCl 0.9%) or anisomycin (10 mg per kg of body mass. Sigma Aldrich) together with the last two saline or LPS administrations. 24 hours after the last injection, mice were euthanized by anaesthesia with ketamine (100 mg per kg of body mass) and xylazine (10 mg per kg of body mass) followed by transcardiac perfusion with PBS. Brains were removed, fixed in cold 4% PFA overnight at 4°C and dehydrated by sucrose infiltration (15% in PBS overnight, followed by 30% in PBS) before processing.

### Immunocytochemistry

Microglia were fixed for 20 min at 4 °C in 4% PFA, 4% sucrose in PBS. Cells were washed three times for 5 minutes with PBS, permeabilized and blocked for 30min in 3% BSA, 100 mM glycine and 0.25% Triton X-100 (all from Thermo Fisher Scientific). Cells were incubated overnight at 4 °C with primary antibodies against puromycin (mouse monoclonal 1:500, Merck), Par3 (rabbit polyclonal 1:1000, Sigma Aldrich), β-actin (rabbit polyclonal 1:200, Sigma Aldrich), IMP1/ZBP1 (mouse monoclonal 1:500, MBL, Woburn MA, USA or rabbit monoclonal 1:50, Abcam, Cambridge, UK), calreticulin (rabbit polyclonal 1:500, Abcam, Cambridge, UK), pS6 (rabbit monoclonal 1:1000, Cell signaling technology, Danvers, Massachusetts, USA), Iba1 (guinea pig, polyclonal 1:500, Synaptic Systems, Göttingen, Germany) and P2Y12 (rabbit plyclonal 1:1000, Sigma Aldrich). Samples were washed three times for five minutes with PBS and incubated with the corresponding fluorescently labelled secondary antibodies for 1 hour at room temperature (all from Invitrogen, Thermo Fisher Scientific). Cells were washed three times with PBS and, whenever stated, labelled with Alexa Fluor 488-conjugated phalloidin (1:800, Thermo Fisher Scientific) in PBS or Alexa Fluor 647-conjugated phalloidin (1:1000, Abcam) in 1% BSA for 30 minutes at room temperature. Samples were washed three times with PBS and mounted with Prolong Gold Antifade mounting medium with DAPI (Invitrogen, Thermo Fisher Scientific).

### Immunohistochemistry

For immunohistochemistry we typically obtained 40 µm-thick vibratome sagittal sections from brain samples. Briefly, brains from perfused mice were transferred to a solution containing 30% ethylene glycol and 30% sucrose in deionized water overnight. Sections were obtained in a Leica vibratome (Leica, Wetzlar, Germany) and stored at - 20°C until used. Samples were washed three times for 5 minutes with PBS, permeabilized and blocked for 30min in 3 % BSA, 100 mM glycine and 0.25% Triton X-100 (all from Thermo Fisher Scientific). Brain sections were then treated with an anti-mouse IgG (donkey polyclonal 1:1000, Jackson Immunoresearch, West Grove, Pensilvania, USA) for 1 hour at room temperature to block unspecific signal and incubated overnight at 4°C with primary antibodies against pS6 (mouse monoclonal 1:1000, Cell signalling technology), IMP1/ZBP1 (mouse monoclonal 1:500, MBL, Woburn, Massachusetts, USA), GFP (chicken polyclonal 1:1000, Aveslab, Davis, California, USA) and/or P2Y12 (rabbit polyclonal 1:1000, Sigma Aldrich) in 3% BSA containing blocking solution. Samples were washed three times for five minutes with PBS and incubated with the corresponding fluorescently labelled secondary antibodies (all from Invitrogen, Thermo Fisher Scientific) together with (DAPI. 4’, 6-Diamino-2-phenylindole, 5mg/ml, Sigma Aldrich). Sections were washed three times with PBS and mounted with Prolong Gold Antifade mounting medium (Invitrogen, Thermo Fisher Scientific).

### SYTO RNA Select labelling

Cultured cells were washed once with cold PBS, once with 50% methanol in PBS and fixed in cold 100% methanol for 5 minutes. Samples were rehydrated by washing them in 50% methanol in PBS once and in PBS three times at room temperature. Cells were incubated for 20 min at room temperature with 500 nM SYTO RNA Select green fluorescent dye in PBS (Invitrogen). Cells were washed three times with PBS and mounted with Prolong Gold Antifade mounting medium with DAPI (Invitrogen).

### Fluorescent i*n situ* hybridization (FISH)

The FISH protocol was performed as previously described ^33^. Cells were fixed for 20 minutes at 4 ^°^C in 4% PFA, 4% sucrose in PBS. RNA was precipitated by washing cells with 50%, 75% and 100% ethanol, and rehydrated by washing once with 75%, once with 50% ethanol, and twice with PBS. Cells were permeabilized and blocked for 30min in 3% BSA, 100 mM glycine and 0.25% Triton X-100 (all from Thermo Fisher Scientific). For antigen retrieval cells were incubated for ten minutes with 1 µg/ml proteinase K (Thermo Fisher Scientific) at room temperature, washed three times with PBS, fixed for 10 minutes with 4% PFA and 4%sucrose in PBS and washed three times for 5 minutes with 3% BSA, 100 mM glycine and 0.25% Triton X-100.

Cells were incubated with 50 ng of *Actb*, *Par3* or *Gfp* (negative control) targeting riboprobes labelled with digoxigenin in hybridization buffer containing 50% formamide (Sigma), 2X SSC (Sigma), 0.2% BSA, 1mg/ml *E.coli* tRNA (Sigma), 1mg/ml salmon sperm DNA (Fisher Scientific) for 3 hours and 30 minutes at 37 °C. As a negative control green florescent protein targeting probes were used. Cells were washed with 50% formamide in 2X SSC once for 30 minutes at 37 °C, once with 50% formamide in 1X SSC for 30 minutes at 37 °C, three times with 1X SSC for 15 minutes each and three times with TBS containing 0.1% Tween at room temperature for 5 minutes each.

Samples were blocked and permeabilized for 30 minutes in 3% BSA, 0.1% Tween in TBS at room temperature. Cells were incubated overnight at 4 °C with primary antibodies against digoxigenin (mouse monoclonal 1:500, Sigma), β-actin (rabbit polyclonal 1:200, Sigma) and/or Iba1 (guinea pig, polyclonal 1:500, Synaptic Systems, Göttingen, Germany) in 3% BSA, 0.1% Tween in TBS. Cells were washed three times with TBS containing 0.1% Tween at room temperature for 5 minutes each and incubated with the appropriate fluorophore-conjugated secondary antibodies for 1 hour at room temperature. Samples were washed with PBS twice for 5 minutes. Coverslips were airdried and mounted with Prolong Gold Antifade mounting medium with DAPI (Invitrogen, Thermo Fisher).

All the digoxigenin-conjugated riboprobes were synthetized with the T7 promoter (…GCCCTATAGTGAGTCGTATTAC-3’) which were preceded by the following specific mRNA targeting sequences at the 3’ end:

*Actb.1*: 5’-

AACCGTGAAAAGATGACCCAGATCATGTTTGAGACCTTCAACACCCCAGC -3’

*Actb.2*: 5’-

ATGTGGATCAGCAAGCAGGAGTACGATGAGTCCGGCCCCTCCATCGTGCA -3’

*Actb.3*: 5’-

AGACCTCTATGCCAACACAGTGCTGTCTGGTGGCACCACCATGTACCCAG -3’

*Actb.4*: 5’-

GAGCGTGGCTACAGCTTCACCACCACAGCTGAGAGGGAAATCGTGCGTGA -3’

*Actb.5*: 5’-

TCCCTGGAGAAGAGCTATGAGCTGCCTGACGGTCAGGTCATCACTATCGG -3’

*Par3.1*: 5’-

GCCGTGCGGAGATGGCCGCATGAAAGTTTTCAGCCTTATCCAGCAGGCGG -3’

*Par3.2*: 5’-

TTCCACGAGAATGACTGCATTGTGAGGATTAACGATGGAGATCTTCGAAA -3’

*Par3.3*: 5’-

GGACGGTGGGATTCTAGACCTGGATGACATCCTCTGTGACGTTGCCGATG -3’

*Par3.4*: 5’-

TGCTTTTCGGCCTTATCAAACCACAAGTGAAATTGAGGTCACGCCTTCAG -3’

*Par3.5:* 5’-

TGCAGATTTGGGGATCTTCGTTAAGTCCATCATTAACGGGGGAGCTGCAT -3’

*Egfp.1:* 5’-

GATGCCACCTACGGCAAGCTGACCCTGAAGTTCATCTGCACCACCGGCAAG -3’

*Egfp.2*: 5’-

GACCACATGAAGCAGCACGACTTCTTCAAGTCCGCCATGCCCGAAGGCTAG -3’

*Egfp.3*: 5’-

ACTTCAAGGAGGACGGCAACATCCTGGGGCACAAGCTGGAGTACAACTACG -3’

*Egfp.4*: 5’-

AAGCAGAAGAACGGCATCAAGGTGAACTTCAAGATCCGCCACAACATCGAG -3’

*Egfp.5*: 5’-

AGTTCGTGACCGCCGCCGGGATCACTCTCGGCATGGACGAGCTGTACAAGG -3’

A similar FISH protocol was followed in mouse brain sections (cortical samples), except that 33,5 ng/μl of *Actb* or *Par3* targeting riboprobes were used in this case. After anaesthesia with 100 mg/kg ketamine and 10 mg/kg xylazine, 1-month-old mice (male and female) were transcardially perfused with NaCl 0.9%. Brains were removed and fixed in cold 4% PFA in PBS overnight at 4°C. Samples were dehydrated first in 15% sucrose in PB 0.1M and later in 30% sucrose in PB 0.1M, cryopreserved in OCT (Thermo Fisher Scientific), and 10 µm-think serial sagittal sections were obtained in a cryostat (Leica, Wetzlar, Germany). Sections were stored at -80 °C until used. Brain samples were co-stained with antibodies against GFP (chicken polyclonal 1:1000, Aveslab) and P2Y12 (rabbit polyclonal 1:1000, Sigma Aldrich) in this case, ad counterstained with DAPI.

In the case of *Actb* detection in hippocampal microglia we used vibratome processed sections, given that 10 µm-thick samples were not appropriate to visualize phagocytic pouches.

### Puromycilation coupled with proximity ligation assays (PuroPLA)

Cells were treated with 150 ng/ml LPS (Sigma) or vehicle as previously described and exposed to 2μM puromycin (Sigma) for 2-10 minutes. Samples were fixed in 4% PFA and 4% sucrose in PBS, washed three times with PBS for 5 minutes and permeabilized and blocked for 30 min in 3% BSA, 100 mM glycine and 0.25% Triton X-100 (all from Thermo Fisher Scientific). Cells were incubated overnight at 4 °C with an anti-puromycin antibody (mouse monoclonal 1:500, Merck) combined with an anti-β-actin antibody (rabbit polyclonal 1:1000, 07-Sigma) or an anti-Par3 antibody (rabbit polyclonal 1:200, Sigma). Detection of newly synthesized β-actin or Par3 through PuroPLA was performed according to the manufacturer’s instructions, using rabbit PLAplus and mouse PLAminus probes and red Duolink detection reagents (Sigma Aldrich). Coverslips were incubated with Alexa Fluor488-conjugated phalloidin (1:800; Thermo Fisher Scientific) for 30 minutes at room temperature, before mounting with with Prolong Gold Antifade mounting medium with DAPI (Invitrogen, Thermo Fisher).

### *Imp1* knockdown experiments

*Imp1* genetic silencing was performed using small interference RNAs (siRNA) in serum-free medium in the absence of antibiotics. Double stranded siRNAs were transfected with lipofectamine RNA iMAX (Invitrogen, Thermo Fisher) following manufacturer recommendations. Cells were transfected for 6 or 24 hours with Imp1-targeting siRNAs (siRNA #1: s155193, or siRNA #2: s155193. Both from Ambion, Thermo Fisher Scientific).

The sequences of *Imp1*-targeting siRNAs are the following:

siRNA #1: 5’ – GGAAAAUACAGAUCCGGAAtt – 3’

siRNA #2: 5’ – GCCUGAGAAUGAGUGGGAAtt – 3’

Transfection efficiency was analyzed by immunocytochemistry with an anti-IMP1/ZBP1 antibody (mouse monoclonal 1:500, MBL).

### RNA sequencing

Microglia were seeded on top of a 1 µm-pore membrane of a transwell system. Cell material from the upper compartment was removed with a cotton swab to isolate PeMPs and viceversa. Total RNA from PeMPs and whole lysates was isolated using the Direct-zol RNA Miniprep kit following manufacturer instructions (ZYMO Research, Irvine, California, USA). RNA quality was addressed in a Bioanalyzer electrophoresis system (Agilent Technologies, Santa Clara, California, USA). After removing rRNAs, 2 ng of RNA was used to prepare total RNA libraries with the SMARTer Stranded Total RNA-Seq Kit (v3, Takara Bio, Kusatsu, Japan). Sequencing was performed in a NovaSeq 6000 instrument (paired-end, 2x 100 bp. Illumina, San Diego, California, USA). Reads were aligned to the rat refence genome (mRatBN7.2) using HISAT2 ^41^ and differential localization to PeMPs was analyzed with DESeq2 ^42^. We then focused on those RNAs with similar or higher levels in PeMPs compared to whole lysates.

### Live cell imaging

To address PeMP motility, microglia obtained from a mixed glial culture were seeded in 35 mm µ-Dish polymer coverslip bottom dishes coated with PDL (90.000-100.000 cells/cm2. Ibidi). Cells were treated with 150 ng/ml LPS or PBS just before live imaging was performed. Three random fields per sample were imaged for 2 minutes with 5 second intervals. PeMP motility was addressed in cells immediately treated with vehicle or LPS and after 5, 10, 15, 20 and 30 minutes of treatment. PeMP motility was measured as the absolute movement of lamellae from a starting position (specified at t=0) regardless of their protrusion or retraction. Average protrusion in cells treated for 10 minutes is also specified in the Results section.

### Transwell migration assays

Microglia were obtained from a mixed glial culture as previously described. Cells (60.000-80.000 cells/cm^2^) were seeded in a transwell system which consists of a culture insert with a polyethylene terephthalate membrane that creates two compartments. Cells were seeded onto PDL-coated membranes with 3 µm-pores which enabled microglial migration toward the lower compartment, or with 1 μm-pores in which cell body migration was restricted. The lower compartment was treated with 150 ng/ml LPS or PBS for 30 minutes. Migration of the nuclei and the cytoskeleton were addressed by DAPI and phalloidin staining respectively.

### In vitro phagocytosis assays

The protocol for *in vitro* phagocytosis was adapted from Beccari and colleagues ^43^. Briefly, SH-SY5Y cell lines stably transfected with the red fluorophore tFP602 (SH-SY5Y vampire. InnoProt, Derio, Spain) were cultured in DMEM (Gibco, Thermo Fisher Scientific) containing 10% fetal bovine serum (Sigma Aldrich), penicillin/streptomycin/glutamine (Gibco, Thermo Fisher Scientific) and 250 μg/ml geneticin (Gibco, Thermo Fisher Scientific). Apoptosis was induced by adding 3 µM staurosporine (Sigma Aldrich) for 4 hours at 37 °C. The medium was collected and centrifuged for 4 minutes at 1300 rpm. Apoptotic cells were resuspended in microglia plating medium and added to microglial primary cultures in a 1:3 ratio (SH-SY5Y:microglia). Primary microglia were exposed to LPS or vehicle for 30 minutes prior to adding the apoptotic cells. Phagocytosis was addressed by quantifying the percentage of phalloidin-labelled microglia containing vampire particles as well as the particle size.

### Image acquisition and fluorescence quantification

Images were acquired in cultured microglia using an EC Plan-Neofluar 40×/1,30 Oil DIC M27 objective on an Axio-Observer Z1 microscope equipped with AxioCam MRm Rev. 3 (Zeiss, Oberkochen, Germany) and Hamamatsu EM-CCD ImagEM (Hamamatsu Photonics, Hamamatsu, Japan) digital cameras. Settings for image acquisition were determined in a random field of a control sample ensuring pixel intensities were within the linear range and avoiding pixel saturation. Images were acquired with ZEN 2 (blue edition) version 2.0.0.0. software (Zeiss). Cells were selected based on the counterstain (e.g phalloidin) for the blind acquisition of the labelling of interest and settings were kept identical for all sampled cells in any given experiment. Whenever possible, five random fields per coverslip and two coverslips per experimental condition were imaged.

In the case of microglia sampled from cryostat-processed mouse brains, images were acquired using the same EC Plan-Neofluar 20x objective microscope in cryostat slices. GFP-positive cells were randomly imaged from the IV layer of the cortex of 10 µm-thick brain sagittal sections. 6 stacks were imaged per field using structured illumination with Apotome2 (Zeiss) and the summed mean pixel intensity was used for quantification. 6 stacks were imaged per field using structured illumination with Apotome2 (Zeiss) and the summed mean pixel intensity was used for quantification.

Images from vibratome slices were obtained in a confocal microscope (Leica Stellaris 5, DM6 B CS. Leica). GFP-positive cells were randomly imaged from the layer IV of the or the dentate gyrus. Individual GFP+ microglia were imaged with a 40X objective, with bidirectional scanning and optical sections of 0.7 μm z-step interval.

In any case, after background subtraction, the average pixel intensity was calculated for each sample. For presentation in figures the greyscale images were converted to RGB and contrast and background settings were set the same in control and experimental conditions for the staining of interest. Markers used as counterstain for PeMP selection were adjusted for an optimal visualization in figures.

SYTO-, FISH- and IMP1/ZBP1-positive foci were measured on binarized images as previously described ^29^. The number of objects was scored for each staining in MPPs or the soma with the Analyze Particle function in FIJI/ImageJ (NIH) and normalized to area or MPP length. Given the discrete numbers of Puro-PLA puncta these were manually quantified on raw images.

### Statistical analysis

The sample size is specified in the figure legends. Statistical analyses were performed with Prism 8 and 10 (GraphPad Software, San Diego, CA, United States) following a randomized block design where samples from the same experiment were matched to eliminate inter-experimental variability, unless matching could not be performed or was statistically not significant. When comparing the means of two groups taking one variable into account, two-tailed t-tests were performed. One-way ANOVA followed by Holm-Sidak’s multiple comparison test was used if one variable was compared between more than two groups, unless otherwise stated. If more than one variable was analyzed, we performed two-way ANOVA followed by Holm Sidak’s or Tukeýs multiple comparison tests.

## ACKNOWLEDGEMENTS

We thank Dr. Ana Maria Aransay from the CICbioGUNE Genome Analysis Platform for her assistance in library preparation and RNA-Seq experiments. We thank Dr. Federico Soria for his attempt in infecting microglia with adenoviruses *in vivo*. We thank Dr. Mariana Astiz for critical review of our manuscript and for her suggestions to improve our work.

## FUNDING

This work was supported by grant PID2019-110721RB-I00 funded by MICIU/AEI/10.13039/501100011033 (to JB), a start-up grant from the Ramón y Cajal program (RYC-2016-19837 to JB) and start-up funds from the Basque Foundation for Science (IKERBASQUE to JB). JI-I is a predoctoral fellow funded by the Basque Government. MB-U is a predoctoral fellow funded by the University of the Basque Country (UPV/EHU).

## AUTHOR CONTRIBUTIONS

JB conceived the project and designed the experiments together with JI-I and MB-U. JI-I and MB-U performed and analyzed the experiments together with JB. IN-G prepared all samples for *in vivo* analyses. IG-T, LCF-B and SC performed RNA-Seq analyses. MM and AS assisted with phagocytosis assays *in vitro* and prepared SH-SY5Y vampire neuronal cell lines. JI-I and AS identified phagocytic pouches *in vivo*. JB and JI-I drafted the manuscript and prepared the figures. JB, JI-I and AS edited the manuscript.

## DATA AVAILABILITY

The datasets generated for this study are available on request to the corresponding author.

## CONFLICT OF INTEREST

The authors declare no conflict of interests.

## SUPPLEMENTARY FIGURES

**Supplementary figure 1.**
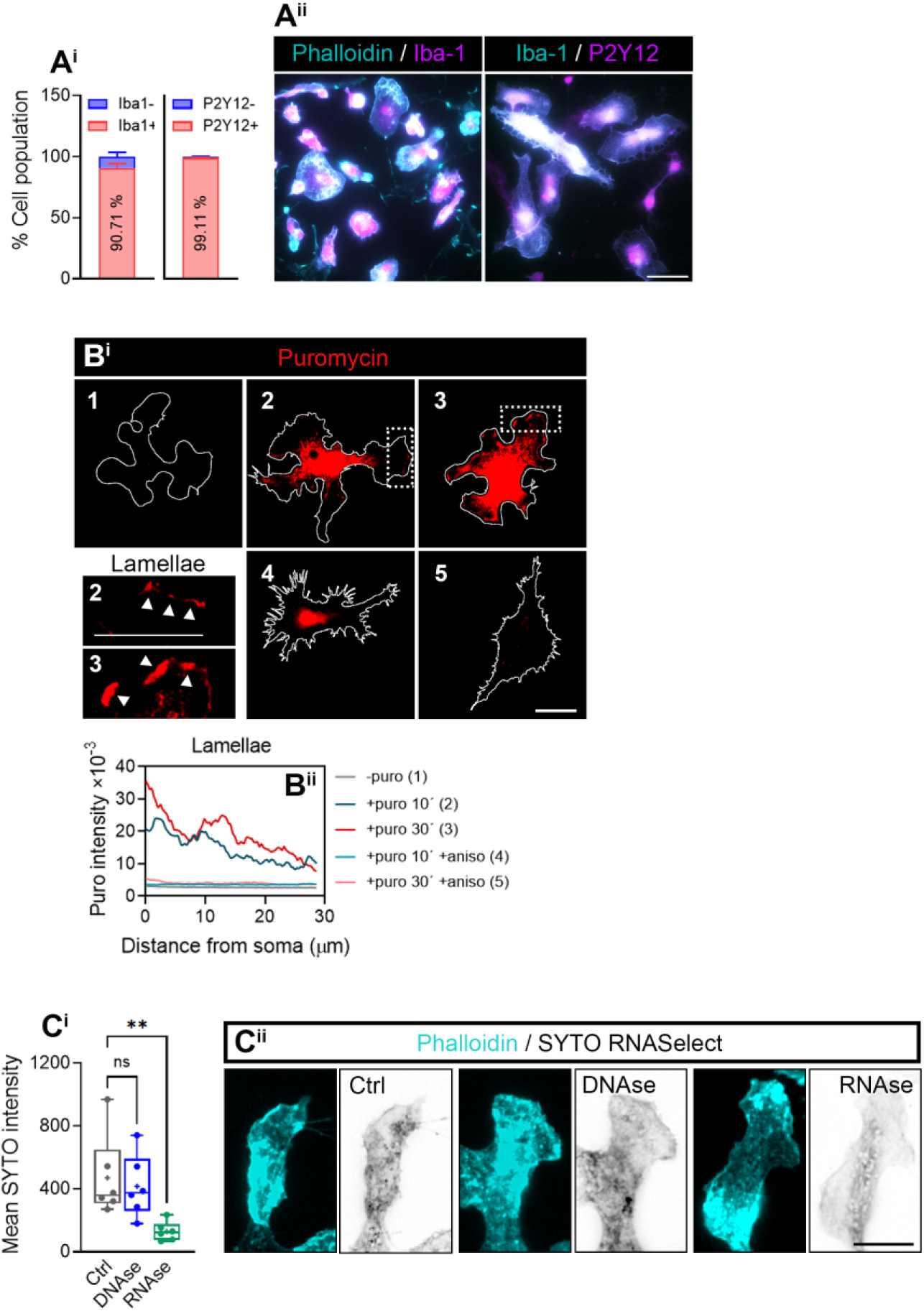
Characterization of primary microglia cultures and methods for detection of newly synthesized proteins and of endogenous RNA in PeMPs. **(A)** Expression of microglia markers in primary cultures. The graphs represent the proportion of phalloidin-positive cells expressing Iba-1 (left) and the percentage of P2P12-positive cells among the Iba-1-positive population. Labelling was performed in cells from 3 independent cultures (n=3) (A^i^). Micrographs show primary cultures stained for phalloidin, Iba-1 and P2Y12. Scale bar 50 µm. **(B)** To visualize protein synthesis in microglial peripheral structures, cells were incubated with vehicle (-puro; 1), with puromycin for 10 and 30 minutes (+ puro; 2 and 3 respectively) or with anisomycin for 30 minutes and puromycin for 10 (+ puro 10’ + aniso; 4) or 30 minutes (+ puro 30 ’ + aniso; 5). Representative micrographs of puromycin distribution in whole cells and in lamellae are shown. Scale bar, 20 µm (insets, 5µm) (B^i^). The graph represents the distribution of puromycin-labelled newly synthesized proteins along lamellae. **(C)** RNA localization to the periphery of microglia was visualized with SYTO RNASelect green fluorescent dye. To determine if SYTO selectively labelled RNA, fixed cells were treated with DNAse or with RNAse. The box and whisker graph indicates the mean fluorescence intensity of SYTO in phalloidin-labelled microglia from 6 independent experiments (n=6) analyzed by one-way ANOVA followed by Dunnet’s multiple comparison test. **p < 0.01; ns, not significant (C^i^). Micrographs show phalloidin and SYTO labelling in control, DNAse- and RNAse-treated cells. Scale bar, 10 µm (C^ii^).

**Supplementary figure 2.**
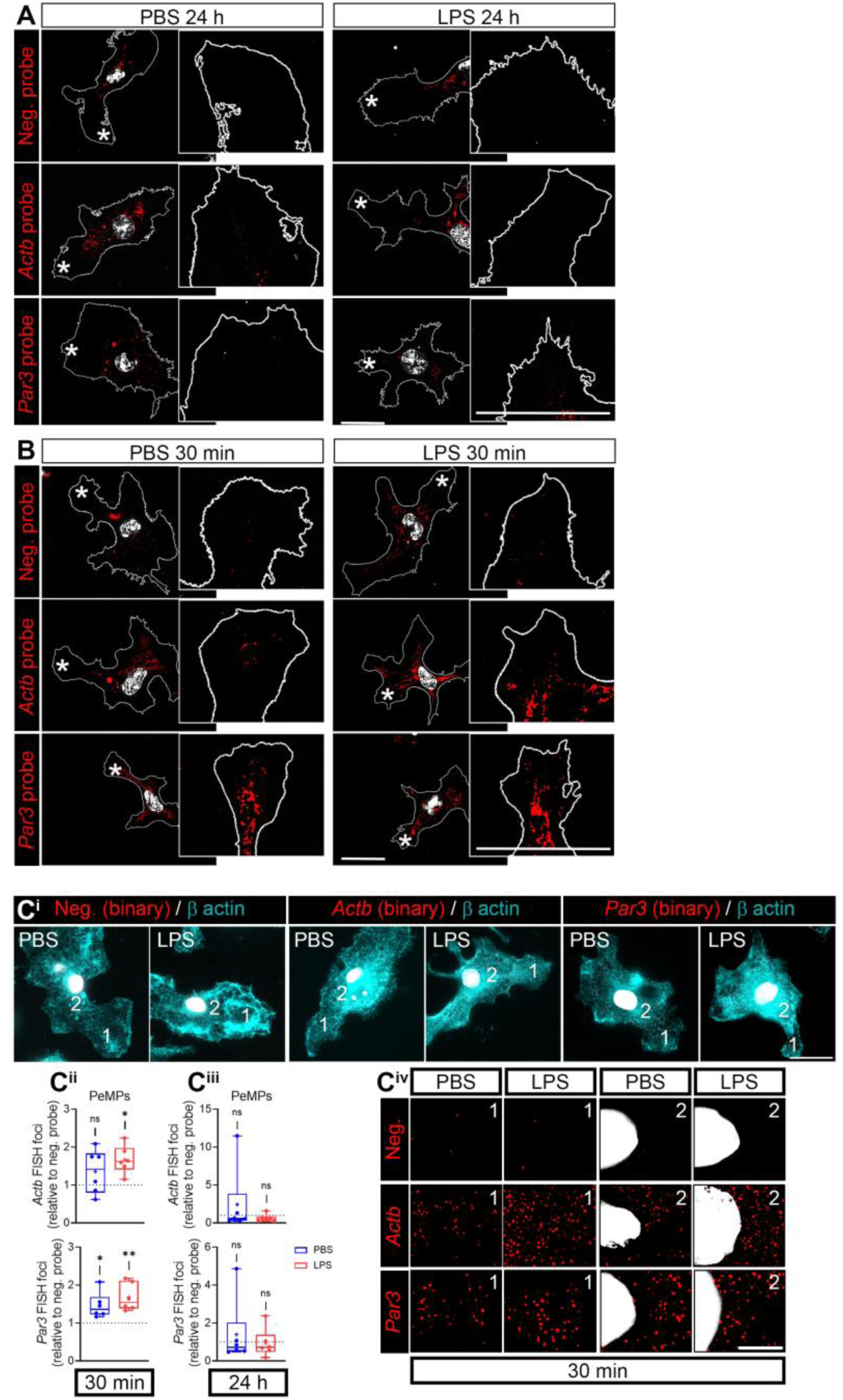
*Actb* and *Par3* mRNA levels in microglia upon LPS exposure. **(A)** A prerequisite for localized translation is the presence of mRNAs in cell peripheral structures. We addressed the localization of *Actb* and *Par3* mRNAs to microglial lamella by fluorescent *in situ* hybridization (FISH) and did not observe any positive signal compared to a negative when cells were exposed to PBS or LPS for 24 hours. Representative images of FISH are shown. Scale bar, 20 µm. **(B)** Both Actb and Par3 were readily detected in cells treated with vehicle or LPS for 30 minutes. Representative images are shown. Scale bar, 20 µm. Asterisks in **(A)** and **(B)** indicate the lamellae shown in insets. **(C)** Binarized FISH signals in cells counterstained for β actin. (1) and (2) in (C^i^) indicate the PeMPs and the perinuclear region represented in insets in (C^iv^). Scale bars 20 µm in (C^i^) and 10 µm in (C^iv^). These are the same cells shown in Figures 2B (for *Actb*) and S3B (for Par3) without counterstain. The box and whisker plots indicate the number of *Actb* (upper graph) and *Par3* (lower graph) mRNA foci relative to the negative probe quantified in 6 independent experiments (n=6) following a 30-minute exposure to PBS and LPS. One-way ANOVA followed by Dunnet’s multiple comparison test. *p < 0.05; **p <0.01; ns, not significant (C^ii^). The box and whisker plots indicate the number of *Actb* (upper graph) and *Par3* (lower graph) mRNA foci relative to the negative probe quantified in 7 independent experiments (n=7) following a 24-hour exposure to PBS and LPS. One-way ANOVA followed by Dunnet’s multiple comparison test. ns, not significant (C^iii^).

**Supplementary figure 3.**
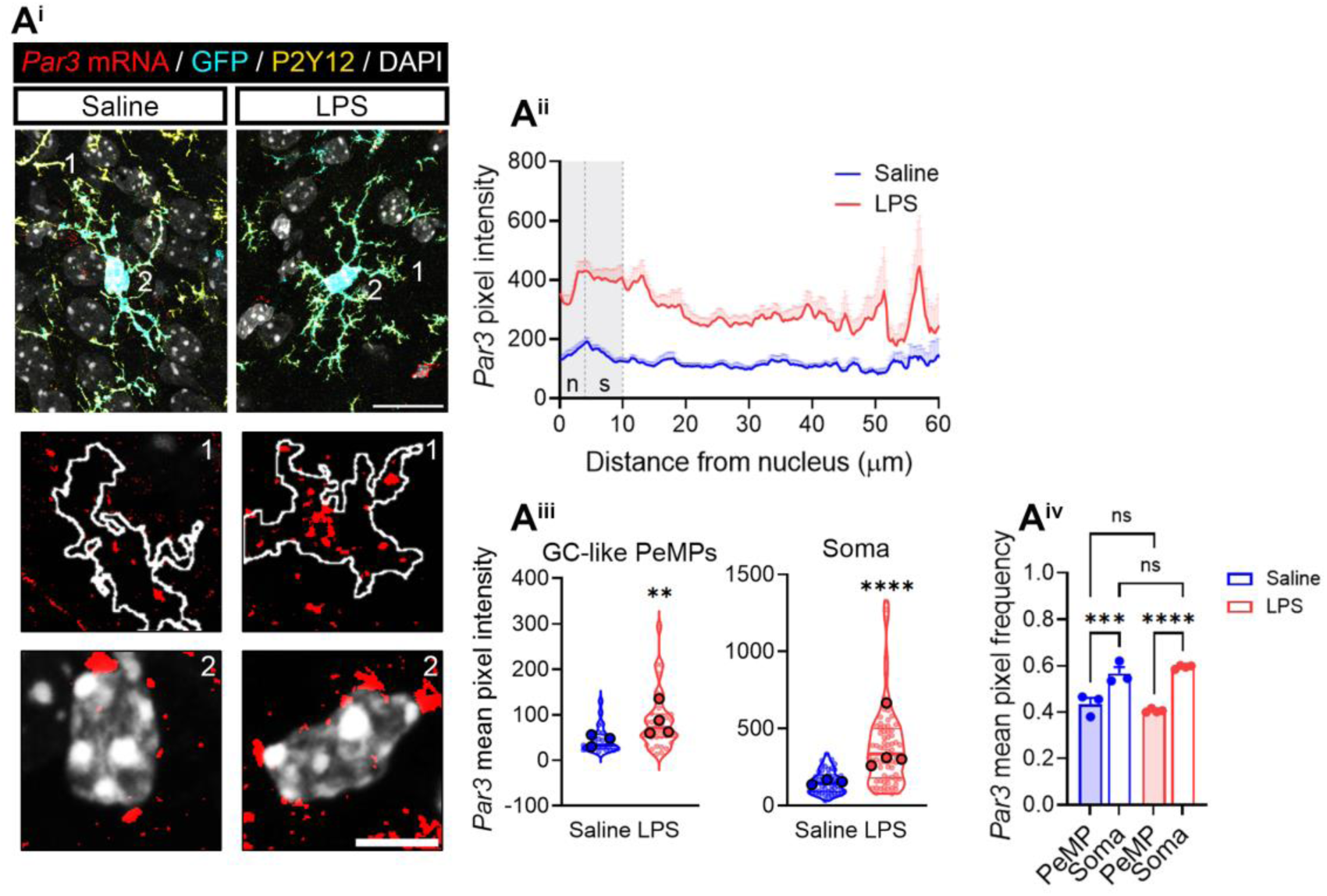
*Par3* mRNA in PeMPs *in vivo*. **(A)** *Par3* levels measured by FISH in cortical microglia (positive for P2Y2 and GFP) from fms-EGFP 1-month old mice injected with saline or LPS. (1) and (2) indicate GC-like PePMs and cell bodies represented in insets. Scale bars 10 µm (5 µm insets) (A^i^). Linescans represent the distribution of the FISH signal from the nucleus to the PeMPs in 49-72 individual cells per condition (A^ii^). Violin plots represent the mean intensity of *Actb* in 25-36 sampled GC-like PeMPs (smaller dots) and 49-72 cell bodies (smaller dots) from 3-4 mice (larger dots). Two-tailed t tests. **p < 0.01; ****p < 0.0001 (A^iii^). The box and whisker graph shows the relative frequency distribution of *Par3* intensity in PeMPs and the soma in cortical microglia from 3-4 mice (n=3-4). One-way ANOVA followed by Holm-Šídák’s multiple comparison test. ***p < 0.001; ****p > 0.0001 (A^iv^).

**Supplementary figure 4.**
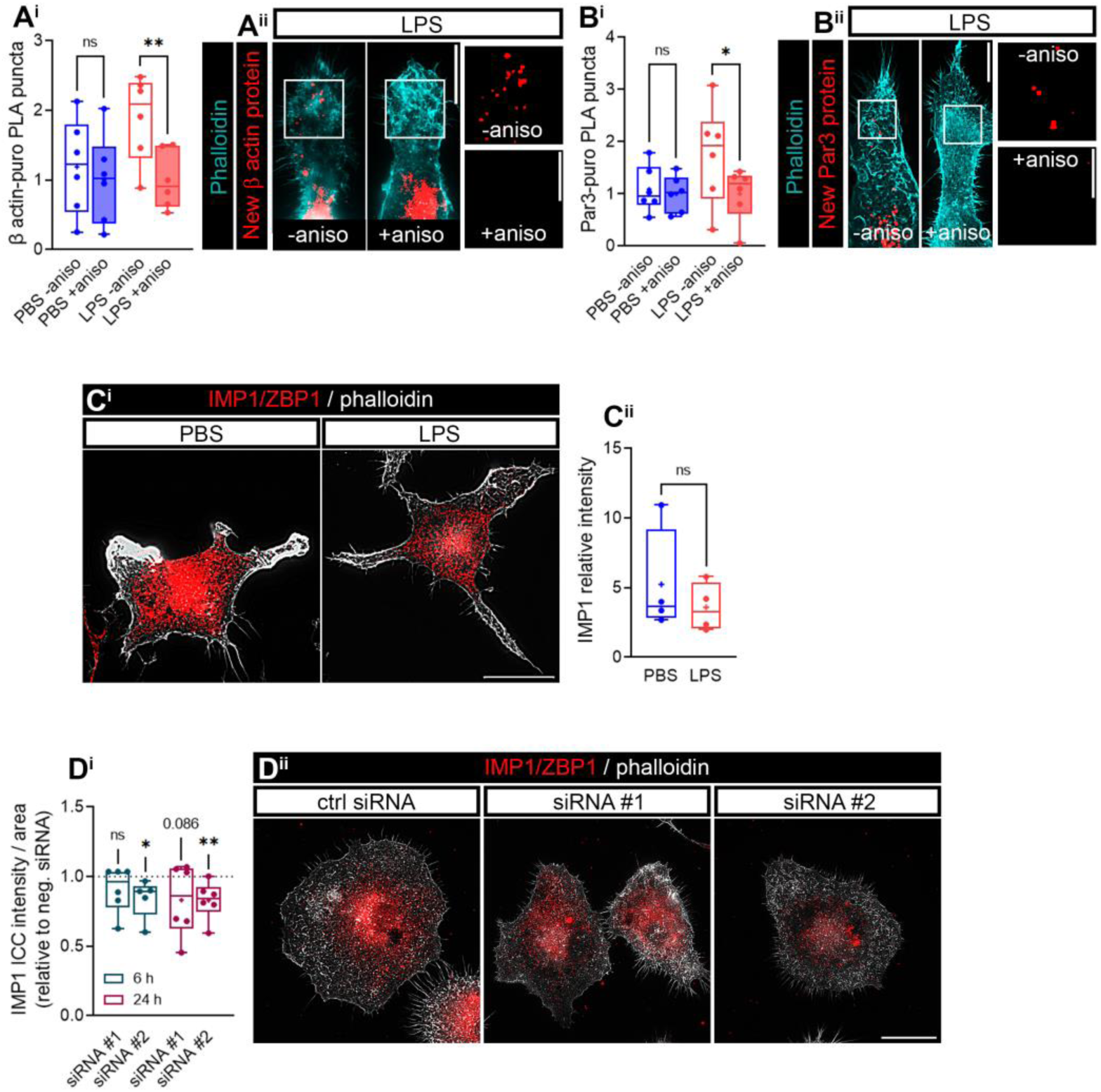
LPS enhances localized *Actb* and *Par3* translation in the periphery of microglia. Analyses of RBP IMP1/ZBP1 in microglia in unstrasfected and transfected cells. **(A)** New synthesis of β actin in microglial peripheral structures was assessed with a 10-minute puromycin pulse in PBS- and LPS-treated cells followed by proximity ligation assay (PLA) with antibodies against puromycin and β actin. As a negative control, cells were preincubated with the protein synthesis inhibitor anisomycin. The box and whisker graph represents the average PLA puncta within lamellae in phalloidin-stained microglia treated with PBS or LPS in 6 independent cultures (n=6) and analyzed by pairwise comparisons with Holm-Šídák’s correction. *p < 0.05; ns, not significant (A^i^). Representative micrographs of PLA puncta in LPS-treated cells are shown. Scale bar, 10 µm (insets, 5 µm) (A^ii^). **(B)** New synthesis of Par3 in PeMPs was assessed with a 10-minute puromycin pulse in PBS- and LPS-treated cells followed by PLA with antibodies against puromycin and Par3. As a negative control, cells were preincubated with the protein synthesis inhibitor anisomycin. The box and whisker graph represents the average of PLA puncta within lamellae in phalloidin-stained microglia treated with PBS or LPS in 6 independent cultures (n=6) and analyzed by pairwise comparisons with Holm-Šídák’s correction. *p < 0.05; ns, not significant (B^i^). Representative micrographs of PLA puncta in LPS-treated cells are shown. Scale bar, 10 µm (insets, 5 µm) (B^ii^). **(C)** IMP1/ZBP1 levels in PBS- and LPS-treated microglia. Representative micrographs show IMP1 immunostaining in phalloidin-positive cells treated with vehicle or LPS. Scale bar 20 µm (C^i^) The box and whisker graph represents the relative IMP1 levels in microglia exposed to PBS or LPS for 30 minutes in 4 independent experiments (n=4). Two-tailed t test. ns, not significant (C^ii^). **(D)** Knockdown efficiency of two nonoverlapping siRNAs targeting *Imp1* compared to a negative siRNA following 6 and 24 hours of transfection. The box and whisker graph represents the mean intensity of IMP1 protein in cells transfected with a control siRNAs and two *Imp1* siRNAs from 5-6 independent experiments (n=5-6). Two-tailed t tests was performed, *p < 0.05; **p < 0.01; n.s, not significant, siRNA #1 vs ctrl siRNA (dashed line) and siRNA #2 vs ctrl siRNA (dashed line) (D^i^). Representative micrographs of transfected cells are shown. Scale bar, 20 µm (D^i^).

**Supplementary figure 5.**
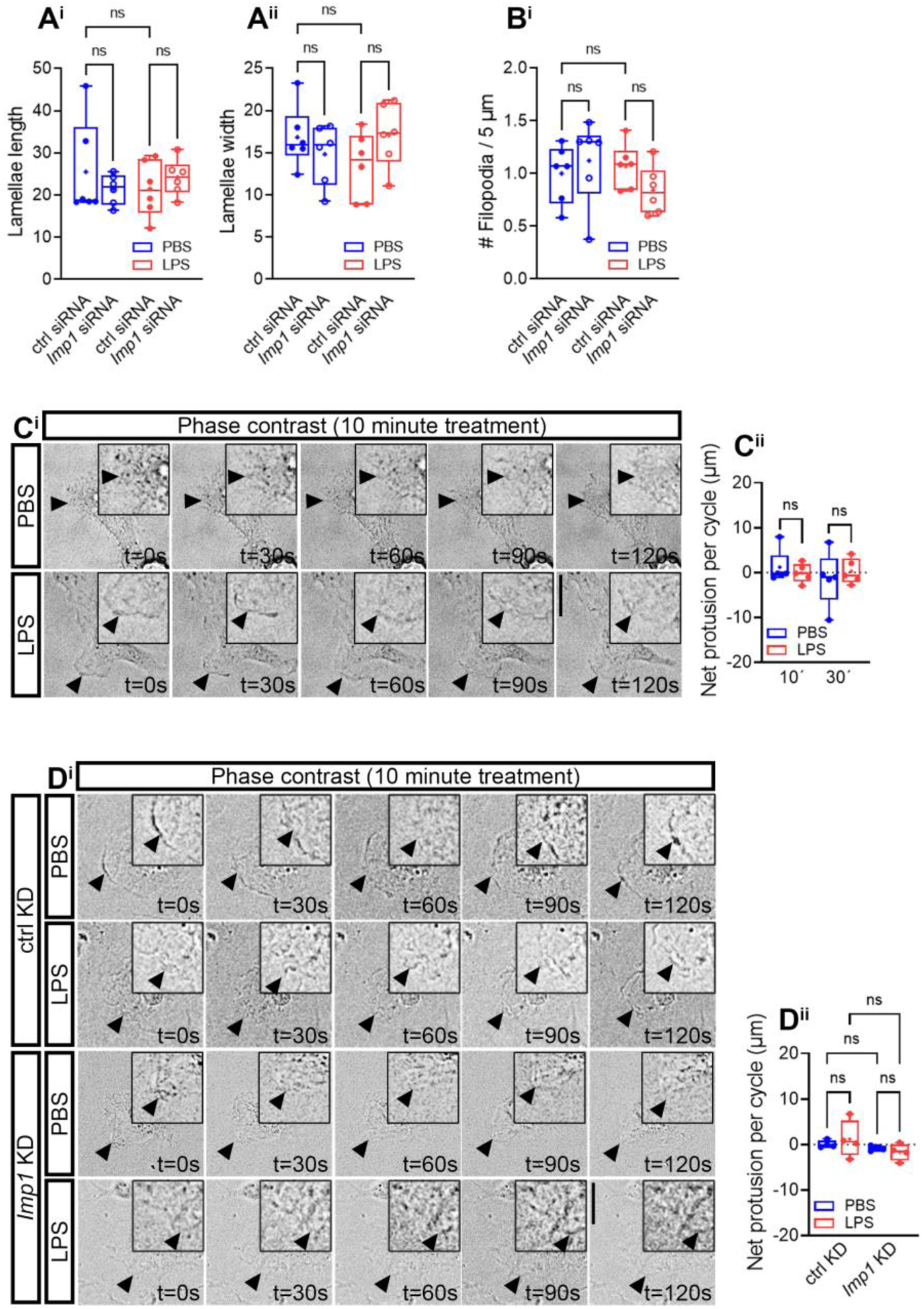
Effect of IMP1/ZBP1 in PeMP morphology and motility in response to acute inflammation. **(A)** Lamellar morphology in PBS- and LPS-treated microglia transfected with a control or an *Imp1*-targeting siRNA. Lamellae length (A^i^) and width (A^i^) were analyzed 6 independent experiments (n=6) as shown in box and whisker plots. One-way ANOVA followed by Holm-Šídák’s multiple comparison test. n.s, not significant. **(B)** Number of filopodia in cells transfected with control (ctrl KD) or *Imp1* siRNA (*Imp1* KD) and exposed to PBS or LPS for 30 minutes. The box and whisker plots represent the results form 6 independent experiments (n=6). One-way ANOVA followed by Holm-Šídák’s multiple comparison test was performed, n.s; not significant. **(C)** PeMP motility analyzed by life cell imaging in untrasfected cells treated with vehicle or LPS 10 minutes. Cell visualization was performed for 2 minutes (5-sec cycles). Representative micrographs of lamellar behaviour every 30 seconds are shown (C^i^). The box and whisker graph represents the net protrusion of lamellae after a 10-minute treatment in cells cultured in 5 independent experiments (n=5). One-way ANOVA followed by Holm-Šídák’s multiple comparison test. n.s, not significant. (C^ii^). **(D)** PeMP motility analyzed by life cell imaging in ctrl KD and *Imp1* KD cells treated with vehicle or LPS 10 minutes. Cell visualization was performed for 2 minutes (5-sec cycles). Representative micrographs of lamellar behaviour every 30 seconds are shown (D^i^). The box and whisker graph represents the net protrusion of lamellae after a 10-minute treatment in cells cultured in 4 independent experiments (n=4). One-way ANOVA followed by Holm-Šídák’s multiple comparison test. n.s, not significant. (D^ii^).

**Supplementary figure 6.**
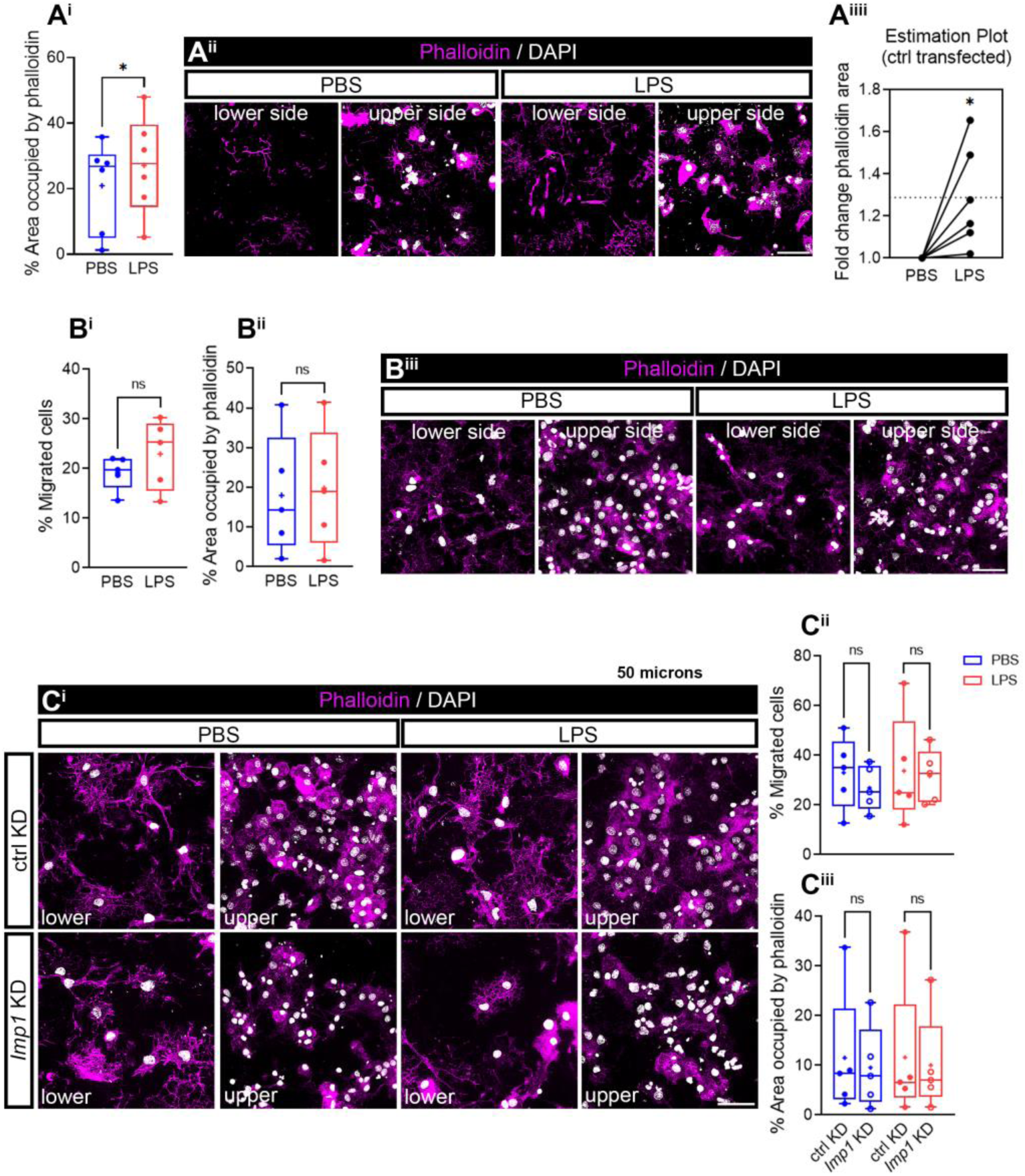
Effect of acute LPS exposure on lamellar polarized migration. **(A)** PeMP migration in a 1 µm-pore transwell culture system. Somatic migration was restricted in this culture setup, and only peripheral processes were observed in the lower chamber. We treated the lower compartment and assessed polarized PeMP migration by the area covered by phalloidin in PBS- and LPS-treated cells. Box and whisker graph represents F-actin-rich cytoskeleton extension (phalloidin staining) measured in 6 independent experiments (n=6). Two tailed t test was performed, *p < 0.05 (A^i^). Micrographs depicting nuclear migration (DAPI) and F-actin extension (phalloidin) toward the lower compartment are shown. Scale bar, 100 µm (A^ii^). LPS effect on cytoskeleton extension was addressed in control- (ctrl) transfected cells to determine the reproducibility of our results compared to those in untransfected cells. Despite the variability, we did observe a consistent relative effect on polarized PeMP migration in LPS-compared to PBS-treated cells. Two tailed t test was performed. p*< 0.05, (A^iii^). **(B)** To address microglial migration, cells were cultured in 3-µm-diameter transwell membranes. PBS or LPS were applied to the bottom of the transwell for 30 minutes. Representative images of cells migrated to the lower side of the membrane and those remaining in the upper compartment are shown. Scale bar, 50 µm (B^i^). The percentage of cells migrated to the lower side of the membrane (B^ii^), as well as the relative coverage of phalloidin staining (B^iii^) were calculated in 5 independent experiments (n=5) and are represented in the box and whisker graphs. One-way ANOVA followed by Holm-Šídák’s multiple comparison test. ns, not significant. **(C)** Microglial cell body migration was also addressed in transfected cells in 3 µm-pore membranes. Representative micrographs of nuclear (DAPI) and F-actin (phalloidin) staining in the lower and upper compartments of transwells is shown. Scale bar, 100 µm (C^i^). Again, cell migration was analyzed as the percentage of microglial nuclei (DAPI) found in the lower side of the chamber with respect to the total nuclei (DAPI) found in both the upper and lower chambers (C^ii^), and the covered by phalloidin in the lower compartment was also measured (C^iii^). Both box and whisker plots represent the results from 5 independent experiments (n=5). Two-tailed t test were performed, n.s; not significant (C^ii^ and C^iii^).

**Supplementary figure 7.**
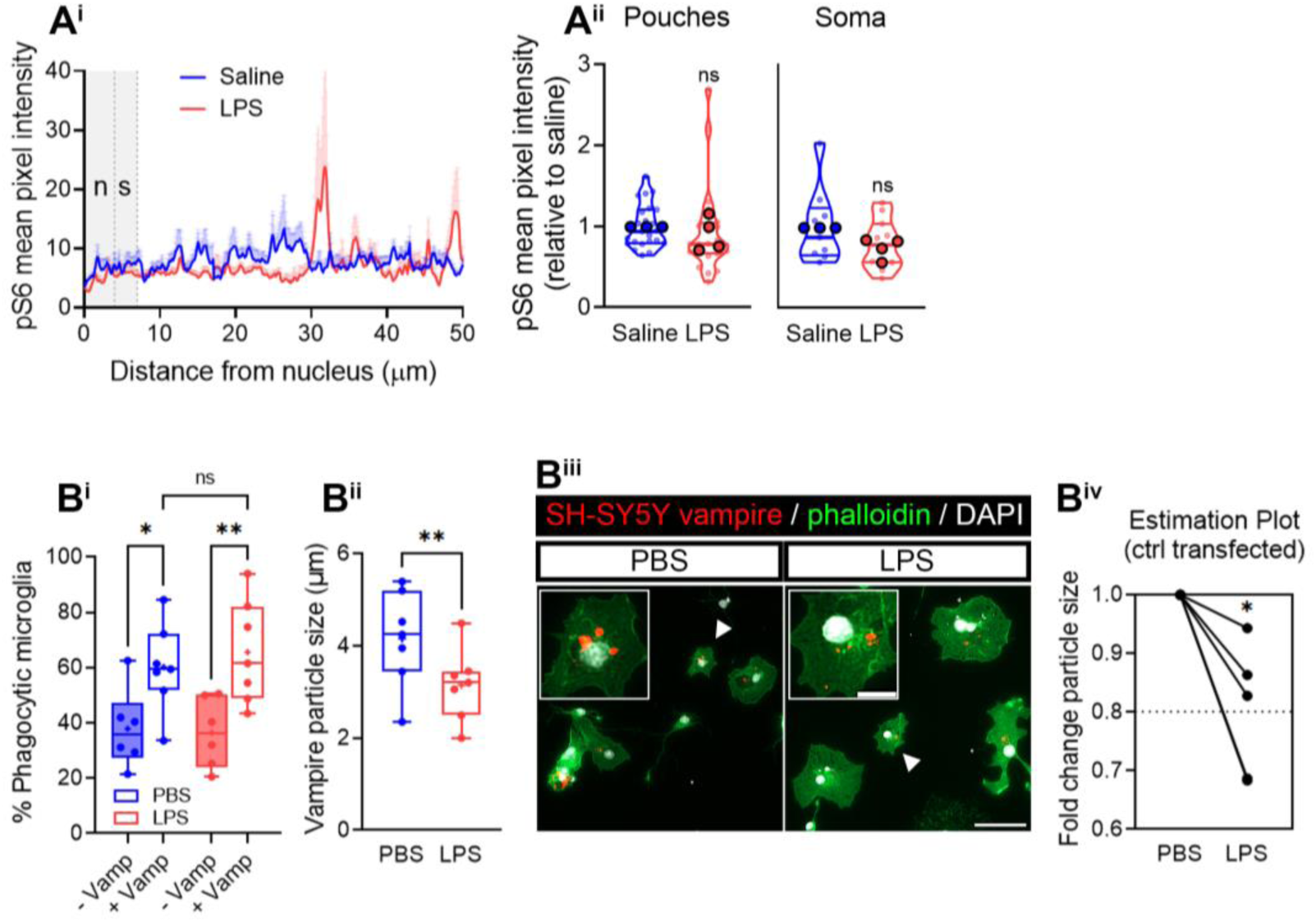
Effect of LPS on phagocytosis. **(A)** Percentage of phagocytic microglia and size of phagocytosed particles in response to LPS. Cultured microglia pretreated with PBS or LPS for 30 minutes were fed for 1 hour with human SH-SY5Y apoptotic neurons labelled with vampire. The percentage of microglia with vampire-positive particles inside was quantified (+Vamp) compared to a negative control (-Vamp). Box and whisker graphs from 7 independent experiments (n=7) show no differences in phagocytic microglia in PBS or LPS treatments (A^i^). However, particles were smaller in LPS-treated cells compared to controls as represented from the box and whisker plots from 7 independent experiments (n=7). Two tailed t test was performed, **p < 0.01 (A^ii^). Vampire particle size in PBS- and LPS-treated microglia is shown in representative micrographs. Scale bar 50 µm (B^iii^). LPS effect on vampire particle size was addressed in control- (ctrl) transfected cells to determine the reproducibility of our results compared to those in untransfected cells. Despite the variability, we did observe a consistent relative effect on the size of vampire particles in phagocytic microglia in LPS-compared to PBS-treated cells. Two tailed t test was performed, *p < 0.05, (A^iv^).

## Notes

### Competing Interest Statement

The authors have declared no competing interest.

### Summary of Updates

This version of the manuscripts has been revised to follow to include suggestions from peers. All figures and a significant part of the text have been edited. Conclusions remain the same.

